# Calibrated Identification of Feature Dependencies in Single-cell Multiomics

**DOI:** 10.1101/2023.11.03.565520

**Authors:** Pierre Boyeau, Stephen Bates, Can Ergen, Michael I. Jordan, Nir Yosef

## Abstract

Data-driven identification of functional relationships between cellular properties is an exciting promise of single-cell genomics, especially given the increasing prevalence of assays for multiomic and spatial transcriptomic analysis. Major challenges include dealing with technical factors that might introduce or obscure dependencies between measurements, handling complex generative processes that require nonlinear modeling, and correctly assessing the statistical significance of discoveries.

`VI-VS` (Variational Inference for Variable Selection) is a comprehensive framework designed to strike a balance between robustness and interpretability. `VI-VS` employs nonlinear generative models to identify conditionally dependent features, all while maintaining control over false discovery rates. These conditional dependencies are more stringent and more likely to represent genuine causal relationships. `VI-VS` is openly available at https://github.com/YosefLab/VIVS, offering a no-compromise solution for identifying relevant feature relationships in multiomic data, advancing our understanding of molecular biology.

## 1 Introduction

Single-cell transcriptomics offers an unprecedented opportunity for probing the function of individual cells and for characterizing the cellular composition of entire samples, thus shedding new light on processes in immunity, development, and pathogenesis of various diseases [1–4]. The emergence of spatial and multiomic technologies further adds the ability to simultaneously profile the surface proteome, epigenome, or location of each cell, on top of its transcriptome. In addition to providing a more comprehensive view of each cell, these technologies open the way for a better understanding of the interplay between molecular or cellular properties. For instance, assessing the dependency between protein abundance on the cell surface and the expression of genes can help identify signaling cascades that help propagate extracellular cues and induce a transcriptional response [5]. Identifying associations between gene expression and the cell’s epigenetic landscape [2] may further help with our understanding of how gene expression is regulated. In spatial transcriptomics, an examination of gene expression patterns across tissue localizations may reveal how the microenvironment affects the function of its residing cells [6]. All of these opportunities require statistical procedures to help detect the most relevant relationships between the observed molecular or cellular features (genes, proteins, chromatin regions, cellularity of the microenvironment, and more).

In single-cell genomics and bulk settings, efforts to detect relationships between such features fall into two broad categories. The simplest methods identify *marginal* associations, which quantify statistical dependencies between pairs of features without considering the other observed features. While these were broadly used for studying gene co-expression networks [7–10], marginal associations suffer from key limitations for single-cell genomics. Practically any technology in this field is impacted by technical factors such as batch effects or variation in sequencing depth, as well as ‘nuisance’ biological factors that are less relevant to the question in hand, e.g., the cell cycle. These factors may inflate marginal correlations, resulting in associations that do not carry the intended biological meaning [11]. More fundamentally, a marginal correlation between two variables in any arbitrary system does not imply causation [12, 13]. For instance, two genes that are regulated by a common set of transcription factors can be highly correlated without being functionally related (Figure 1A). Even when they are functionally related, marginal dependencies may not inform on the proximity of this relationship when two highly correlated genes are indirectly linked through a series of mediator genes (Figure 1B). Marginal approaches hence tend to detect many spurious or indirect associations, which requires further filtering to identify the most relevant relationships [7, 14, 15].

**Figure 1:**
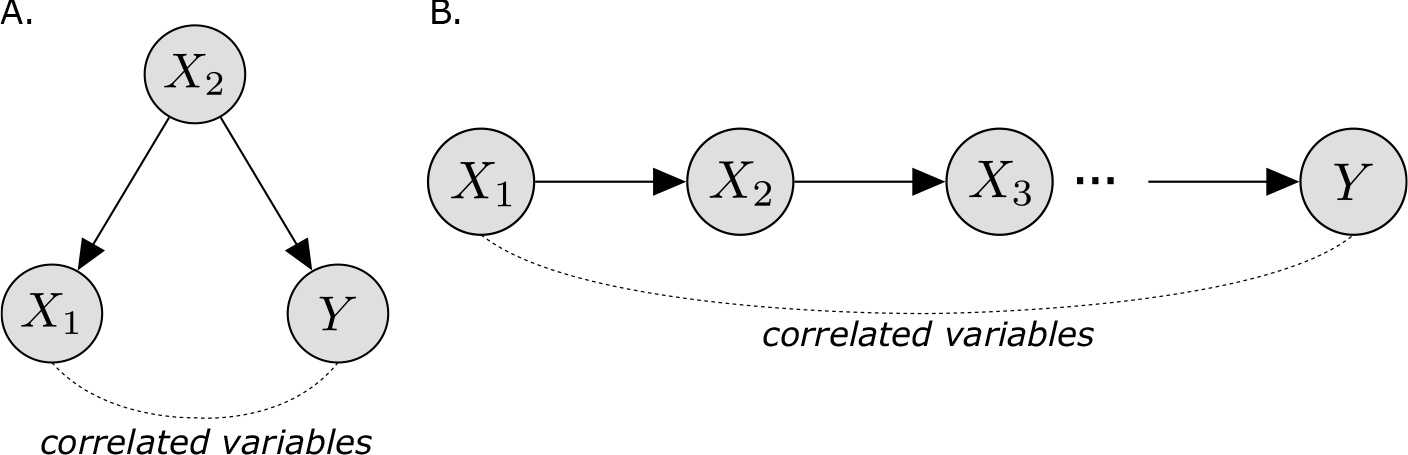
Correlated variables may be functionally unrelated. Here, *X*_1_, *X*_2_, *Y* are random variables characterizing the expression of three genes. **A**. *X*_2_ is directly and causally linked to *Y* and *X*_1_. Here, *X*_1_ and *Y* might be highly correlated, but *X*_1_ does not causally affect *Y* . **B**. *X*_1_ is directly and causally linked to *X*_2_, and similarly, *X*_2_ is connected to *Y* . Here, *X*_1_ and *Y* might be highly correlated, but their association is indirect.

**Figure 2:**
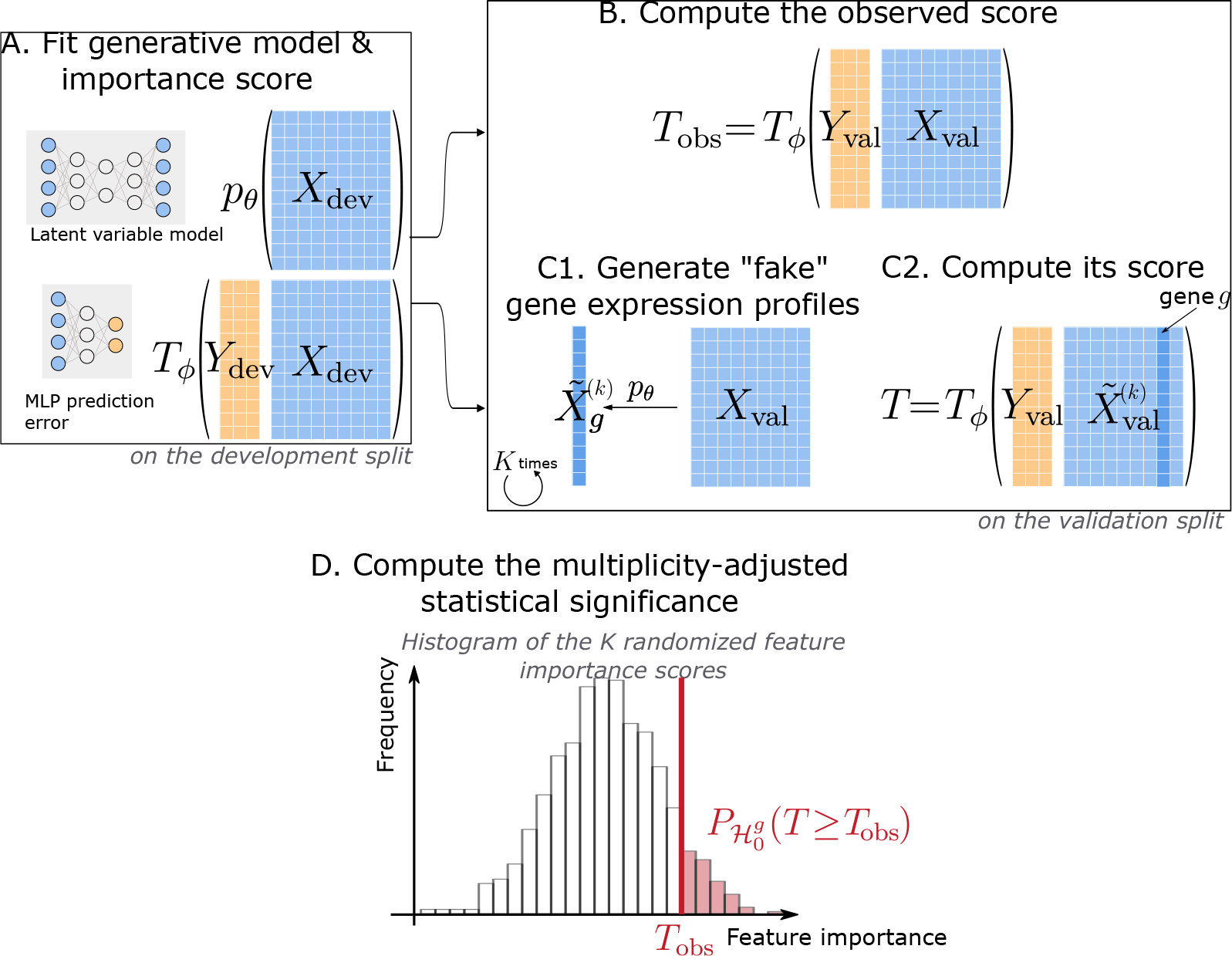
VI-VS pipeline. For simplicity, we omit the confounding factor adjustment from this schematic. **A**. We first randomly split the observed data into a *validation* and a *development* set. We fit a deep generative model *p*_*θ*_ and feature importance score *T*_*ϕ*_, which is a scalar-valued function taking *y* and *x* as inputs on the development set. In a simple case, *T*_*ϕ*_ may correspond to the prediction error of the ordinary least squares of *y* on *x*. Here, *θ* and *ϕ* denote the parameters of these models, learned on the development set. **B**. We compute the feature importance of the observed data. **C**. In parallel, we sample *K* “fake” feature samples for gene *g* using the trained deep generative model (**C1**). Next, we compute the feature statistic of the modified feature matrix *X* where the *g*th column was replaced by generated samples (**C2**). **D**. We rely on the distribution of the “fake” feature statistics to compute the significance of the null hypothesis under which gene *g* and the response are conditionally independent.

*Conditional associations* are a second category of relationships that address these issues by accounting for the overall dependency structure of the data when assessing the dependency between a pair of variables [16–20]. Specifically, detecting conditional dependencies between a response variable *Y* and individual features in a feature matrix *X* often starts by learning a predictor function *f* (*X*) ≈ *Y*, which is then scrutinized to identify variables in *X* that are most associated with *Y* . The simplest example for this approach is the generalized linear models [21, 22], in which learned regression coefficients are used to quantify conditional associations. While limited to linear relationships and simple noise models, linear approaches are relatively scalable. In some cases, these models come with statistical guarantees for the inferred coefficients and are thus easily interpretable. Nonlinear predictors [16, 17] have also been introduced to capture more complex relationships, with tree ensembles being the most prevalent approach. Ensemble approaches have been demonstrated to reach state-of-the-art performance in a variety of tasks such as inference of regulatory interactions between genes [23]. Conditional dependencies therefore provide a more stringent notion of association than marginal dependencies and are more likely to reflect causal relationships. Indeed, pairwise dependencies that persist after conditioning on all other variables imply causal relationships in cases where the causal direction is known, there are no unobserved causal variables, and there is no feedback loops [24]. As such, conditional dependencies are a promising avenue for uncovering relevant interactions in single-cell multiomic data.

In practice, however, algorithms for identifying conditional relationships often need to compromise on (i) *scalability*, e.g., requiring heavy pre-processing to ensure that inference can be completed in a reasonable timeframe [23], (ii) *modeling assumptions*, using often mis-specified view of the underlying process, e.g., with simplified noise models or by assuming linear relationships between variables [25], and importantly: (iii) *interpretability and calibration*, by relying on heuristics to evaluate which of the interactions under consideration are indeed relevant [26–28]. Given these challenges, analyzing dependencies in single multiomics, where millions of measurements (possibly from different batches or studies) are available, requires the use of scalable and rigorous statistical methods. These methods should be able to handle count data distributions, account for technical and biological noise and bias, and allow for nonlinear relationships between variables.

To address these three challenges, we introduce `VI-VS` (Variational Inference for Variable Selection), a general framework for conditional independence testing with multiomic data. `VI-VS` is based on the conditional randomization test (CRT) [29], which quantifies the credibility of pairwise interactions by measuring the effect of exchanging observed features with synthetic ones. We demonstrate and theoretically prove that our procedure provides a calibrated estimation of the false discovery rate. This is achieved without making any assumptions about the distribution of the response variable *Y* or the nature of its interactions with the features in *X*, such as linearity. `VI-VS` harnesses the distributional expressivity of latent variable models, allowing for a variety of noise models for *X*, including count distributions commonly used in single-cell genomics. Finally, `VI-VS` relies on deep neural networks for testing, allowing it to scale to large single-cell genomic datasets as well as capture complex nonlinear relationships between variables.

In the following, we demonstrate the accuracy and calibration of `VI-VS` with several simulation and multi-ome case studies. We also showcase that our procedure provides a theoretically grounded “wrapper” framework that can take existing algorithms for detecting pairwise relationships, and use them to output calibrated decisions. We demonstrate this using the popular GENIE-3 algorithm for inference of regulatory networks.

## 2 The VI-VS model

As an input, `VI-VS` receives a matrix of features *X* ∈ ℝ^*N×G*^ and a vector, representing a response variable *y*∈ ℝ^*N*^ where *G* is the number of features and *N* is the number of cells. We also assume that observed nuisance factors *S* ∈ ℝ^*N×T*^, e.g., batch assignments, sequencing depths, or cell cycle events, affect these experiments and need to be accounted for. Our goal is to detect features in *X* that are associated with the response variable *y* while controlling for the nuisance factors.

In the following, we assume that *X* consists of observed molecular expressions of *G* genes in *N* cells. The choice of *y* varies depending on the assay considered and the problem of interest. Specifically, *y* can characterize molecular quantities, such as protein counts in CITE-seq experiments or chromatin accessibility in ATAC-seq data. It can also represent other, more abstract cell-level properties, e.g., characterizing the tissue environment of a cell in spatial transcriptomic assays. *y* may also correspond to a singled-out gene of interest, for which we wish to identify the interacting genes.

When referring to observations from an individual cell, we will employ lowercase letters, reserving uppercase letters for the entire array of observations. In addition, *x*_*g*_∈ ℕ and *x*_− *g*_ ∈ ℕ^*G*−1^ will respectively denote gene expressions for gene *g* and the vector of remaining genes. When needed, superscripts will index cells, such that *x*^*n*^ denotes the gene expression of cell *n*. When *A* is a set of features, *x*_*A*_ will denote the vector of features contained in *A*. We make the assumption that the samples (*x*^*n*^, *y*^*n*^, *s*^*n*^) are independent and identically distributed (i.i.d.).

### 2.1 Conditional randomization tests for single-cell genomics

To detect genes in *X* that are associated with the response variable *y*, `VI-VS` employs a *conditional* independence test, which estimates, for each gene, the plausibility of the null:

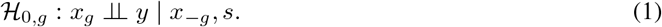

We rely on the conditional randomization test (CRT) approach [29] to test these hypotheses. The premise of CRT is that while it is difficult to directly assess how the distribution of the response variable *y* depends on *X*, it is easier to describe how the features of *X* depend on each other. Leveraging this, our procedure starts by training a predictor function *f* : *x, s ↦y* on held-out data and estimating its reconstruction error. To assess the importance of a feature *x*_*g*_, we *sample* from the distribution of these errors if the null 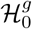 were true. The significance of a feature is determined by comparing the observed error with the error distribution under the null hypothesis. `VI-VS` therefore requires two ingredients: a *generative model* for *X* to capture the dependencies between features, and an *importance score* to evaluate their association with *y*.

#### Importance score

The importance score is a function *T* : *X, Y, S*→ ℝ, which summarizes the observed data. To make decisions, the CRT compares this summary *T* (*X, Y, S*), with 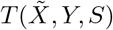), where 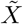 denotes partially synthetic data in which one or few of the features are replaced with values that are generated with the generative model. Here, we propose to *learn* the importance scores from the data. In particular, we consider importance scores corresponding to the prediction error of a regression model of *y* on *X* and *s*,

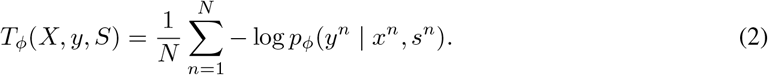

where, *p*_*ϕ*_(*y*_*n*_ | *x*_*n*_, *s*_*n*_) is a likelihood for *y* based on a model *p*_*ϕ*_ trained on held-out data. For instance, *p*_*ϕ*_ may be based on a linear regression, or more complex models such as random forest or a multi-layer perceptron (MLP). Importantly, this predictive model does not need to perfectly capture the conditional distribution of *y* given *X, s*. The CRT will indeed control the false positive rate irrespective of the choice of the predictor model and its assumptions on the nature of the interaction between *X* and *y* or on the distribution of *y* [29]. However, the more adequate the model, the more powerful we can expect the test to be.

#### Generative model

The other required component is a generative model *p*_*θ*_ that (i) can be used to sample “fake” expression for a given gene, and (ii) does not depend on the response variable *y*. Due to their scalability, ability to capture nonlinear effects, and flexible likelihood assumptions, latent variable models are a useful choice to model gene expression in this context. In these models, an unobserved low-dimensional variable *z* is assumed to capture the state of each cell and provide a concise summary of the biological variation among cells. We assume that the model factorizes, for each individual cell and under i.i.d. assumptions, as

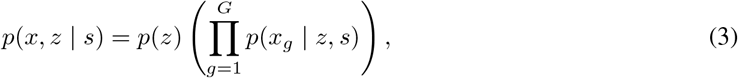

where *p*(*z*) is the latent variable prior, and *p*(*x*_*g*_ | *z, s*) is the likelihood for gene *g*. We rely on variational autoencoders (VAEs) to define the latent variable model. In this model, the prior is usually the standard normal, and the posterior distribution is approximated using a variational approach, with the approximation parameterized by neural networks [30–32]. Assuming access to such a model, testing ℋ_0,*g*_ requires replacing the measurements for the feature *g*, with synthetic measurements that are conditionally independent of *y*. To this end, we use the generative model to obtain K vectors 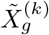, *k* ≤ *K*, containing synthetic counts for gene *g* for all the cells in a manner independent of *y*1. We then construct the overall gene expression for which gene *g* was randomized, as

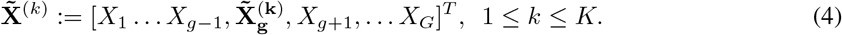

With these two components, a p-value for ℋ_0,*g*_ with the CRT corresponds to the proportion of random trials in which the importance score, when gene *g* is replaced with synthetic data, is not worse than the score obtained with the original data. It writes as

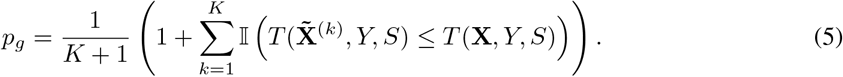

### 2.2 Valid inference for CRTs with latent variable models

Given a latent variable model, an intuitive way to generate synthetic samples 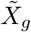 is by independent draws from the Gibbs distribution:

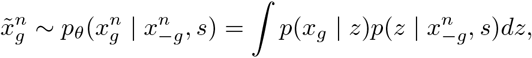

This choice, however, requires sampling from *p*(*z x*_− *g*_, *s*). In the context of VAEs, this requires training a separate model for every feature *g*, which is in most cases computationally prohibitive. Instead, `VI-VS` provides a fast and valid sampling alternative that still provides valid p-values. This is done by drawing *fixed* posterior sample of *z*. Here, for each cell *n*, we first sample one particle from 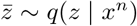 where *q* is the encoder network of the VAE. We then rely on the decoder network of the VAE to obtain synthetic samples:

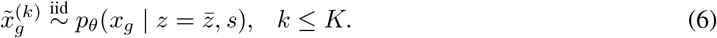

Note that in both cases the generative model does not have access to the value of *y* during sampling, The samples 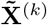 therefore reflect a hypothetical reality in which *x*_*g*_ and *y* are conditionally independent. In Proposition 1 we demonstrate that both sampling schemes provide valid p-values for the CRT (proof in supplemental methods).

#### Proposition 1

(Valid sampling distributions for CRTs with latent variable models). *Assume a latent variable model p*_*θ*_(*x, z* | *s*), *factorizing as* (3). *Let* 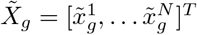 *be a vector of synthetic gene expression profiles generated using the latent variable model for gene g obtained using either of the two sampling schemes described above. Then, the p-values p*_*g*_ *in Equation* (5) *have a distribution that stochastically dominates the uniform distribution when the null hypothesis* ℋ_0,*g*_ *holds. That is, p*_*g*_ *is a valid p-value*.

The entire procedure can therefore be summarized as follows:

#### Algorithm 1 Conditional randomization tests with VI-VS

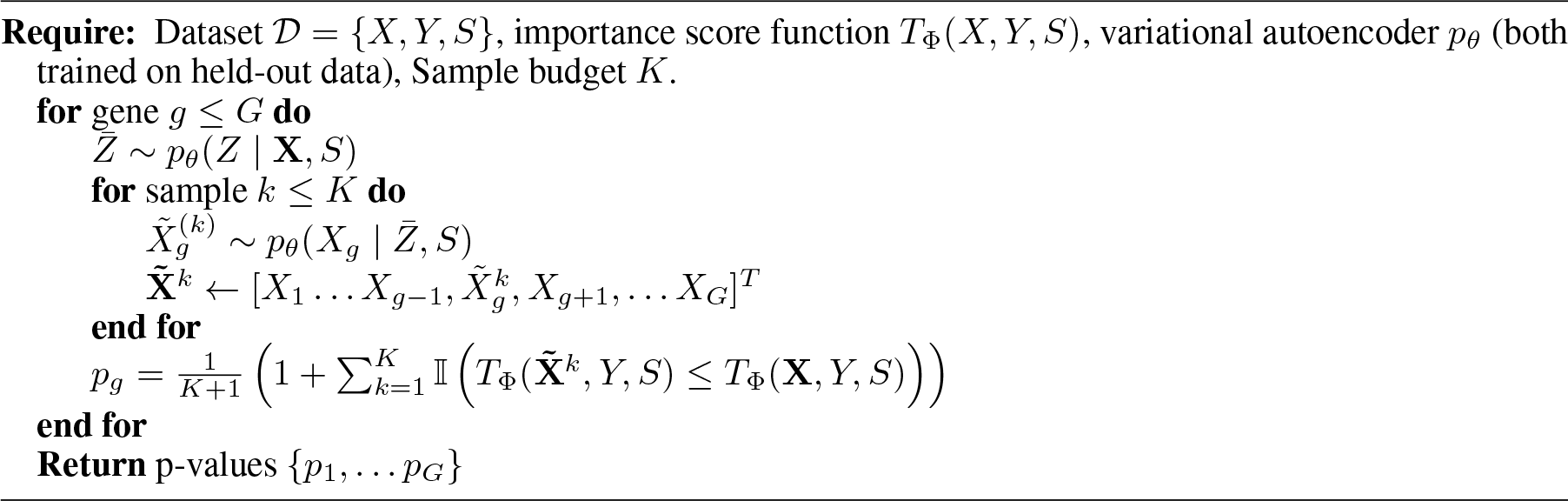

`VI-VS` further corrects the obtained p-values *p*_*g*_ using the Benjamini-Hochberg procedure [33] to control the false discovery rate (FDR), described in Supplement A.1.

#### Cell-specific scores

Equation (5) quantifies the significance of the association between gene *g* and protein *p*; it does not, however, inform on which cell subpopulations may be most responsible for this association. For this purpose, we introduce the cell-specific score,

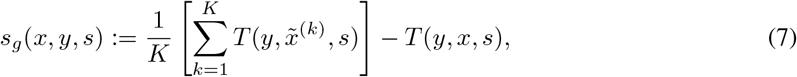

where 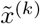 denotes a randomized sample for the CRT. In other words, positive scores will highlight that randomizing the gene *g* in cell *x* decreases the predictive loss, which may mean that this gene plays a relevant role in the prediction *y* for the considered cell.

### 2.3 Multi-resolution hypothesis testing

Conditional dependence is a more stringent statistical notion than marginal dependence. Consequently, applying conditional independence tests at the gene level may not yield many significant genes. This could be due to several factors, such as limited sample sizes or strong correlations between genes that make it challenging to reject the conditional null.

Consider a hypothetical experiment where genes form two clusters and protein expression *Y* conditionally depends on genes 1 and *g* + 1 (Figure 3A). In scenarios with small sample sizes, gene 1 might not be detected if it heavily correlates with other genes in its cluster. However, it is easier to detect that one or more genes in the cluster are conditionally associated with the response, even if we cannot definitively identify which genes are responsible.

**Figure 3:**
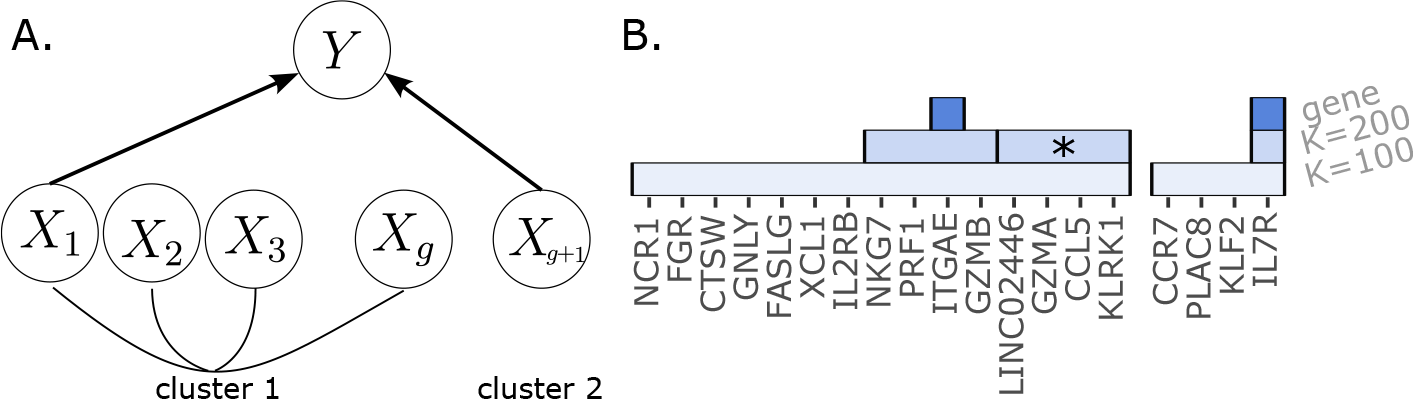
Illustration of the multi-resolution hypothesis testing procedure. **A**. Comparative example of marginal and conditional independence testing. Some surface protein expression causally depends on two genes 1 and *g* + 1. Gene 1 belongs to a cluster of highly correlated genes {1, 2, … *g*) . **B**. Two groups of conditionally dependent genes with the local density of tumor cells for T cells (see Results). Rectangles materialize the groups of genes, or individual genes that are detected by `VI-VS` applied at several resolutions (*K* ∈ {100, 200) and at the gene resolution).

Therefore, following an approach introduced for genome-wide association studies [34], we test for conditional independence at multiple resolutions, ranging from broad groups of genes to the individual gene-level resolution. This strategy helps to avoid overlooking relevant genes such as gene 1 in the above example.

Figure 3B provides a preview of the output of this approach. `VI-VS` identifies two groups of genes at a coarse resolution, as well as significant subgroups at finer resolutions. The first group further breaks down into two significant subgroups at a slightly refined resolution (*K* = 200). In the second group, marked by an asterisk (*), no conditionally dependent gene can be identified. While we cannot pinpoint the specific gene responsible for the association, this analysis suggests that it is due to a biological process distinct from that characterized by ITGAE.

We will now explain how we: (i) group genes together, and (ii) test for conditional independence of a group of genes.

#### Determining relevant clusters of genes

Our goal is to group together features associated with the same biological functions. We assume that high correlations between features may indicate that they are associated with the same biological function. Consequently, we cluster genes based on their empirical correlation matrix. Any gene clustering algorithm can be used in principle, e.g. [35]. We propose using a fast hierarchical clustering approach that is scalable to large datasets. This approach performs agglomerative clustering based on the gene-by-gene empirical correlation matrix computed on the normalized gene expression of the generative model, e.g., scVI [30]. More details about this procedure can be found in Supplement B.1. At a specified resolution *K*, the clustering provides a partition *M* of all genes into *K* groups of genes *A*_1_, …, *A*_*K*_.

#### Group conditional independence

Next, we aim to determine whether a cluster of genes is significant. To formalize this, let *A* ∈ *M* denote a group of genes. We are interested in interactions of the form

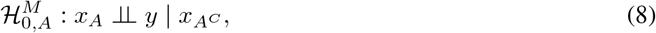

where *x*_*A*_, 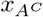 denote the gene expression vectors for the genes in sets *A* and its complement *A*^*C*^, respectively. We test this null hypothesis using the same procedure as described above, with more details provided in Supplement B.1 and illustrated in Figure S1. Specifically, we can test Equation 8 by sampling from the same distribution as in Algorithm 1.

### 2.4 Faster inference using parallel computing

We implemented `VI-VS` in a fast and scalable way that is available as an open-source Python package2. The scalability of this solution relies on two components. Our implementation first relies on parallel computing and just-in-time compilation components of Jax to speed up the inference, allowing us to efficiently compute the p-values for all genes. This practical choice offered a two-fold speedup compared to a Pytorch backend (Supplement B.2). The second key component is the fact that we have set up our algorithm to avoid fitting a model for each Monte Carlo sample, and instead, we only have to perform a forward pass of the pre-fit feature statistic. This computational ingredient improves the run time by orders of magnitude, thus improving scalability.

## 3 Results

### 3.1 Setup

We used scVI [30] as the generative model for gene expression, which was reimplemented in Jax with its default parameters. Importance scores were calculated as the prediction errors of regression models of *y* given *x, s*, either corresponding to a linear model or an MLP. To train these models, we randomly split the available data into a 70%-30% development-validation split. Both the generative model and the importance scores were trained on the development split; p-values and cell scores were computed on the validation split. Note that the generative model and importance scores need only be fit once, upstream of the CRT. In cases where our experiments contained multidimensional response variable *y*, we applied `VI-VS` separately and in parallel to each dimension. In such cases, however, the generative model only needs to be trained once (Supplement B.3).

We compared `VI-VS` with the different importance scores to two baseline approaches. We first considered linear regression of the response variable *y* on the gene expressions matrix *X*, using an Ordinary Least Squares (OLS) objective, representing a baseline method for conditional Independence. We identified conditional associations between genes in *X* and the response *y* by performing a t-test on the corresponding regression coefficients. To study the merit of the conditional independence tests, we also considered a simpler marginal independence test baseline. For each gene *g*, we regressed *y* on the expression of *g* and used a t-test to estimate significance. In both baselines, we used all available data, i.e., both the development and validation splits, to fit the regression models, thus lending a natural advantage over the way `VI-VS` was fit.

### 3.2 VI-VS provides calibrated decisions in a semi-synthetic experiment

We considered a scRNA-seq dataset of 6,855 peripheral blood mono-nuclear cells (PBMC) [36] from a healthy human donor, with 500 genes. We generated five synthetic response variables, each corresponding to the expression of a surface protein measured in the observed cells. The expectations of each response variable were calculated as a linear combination of the squared values of mean-centered log count per million (CPM) expression levels of 150 randomly selected genes in *X*. These simulated response variables were further corrupted by the addition of Gaussian noise. This simulation represents a case where simple linear assumptions do not hold, but there is still a relatively simple model that connects *y* to a subset of features in *X*. More details on data generation can be found in Supplement C.2.

We first evaluated the extent of type I error of the different algorithms (Figure 4A). We found that the FDR estimates of the marginal independence test are poorly calibrated, leading to many false positives. These false positives likely reflect indirect correlations, that is, genes that were not used to generate the synthetic response variable but strongly correlate with other genes that are.

**Figure 4:**
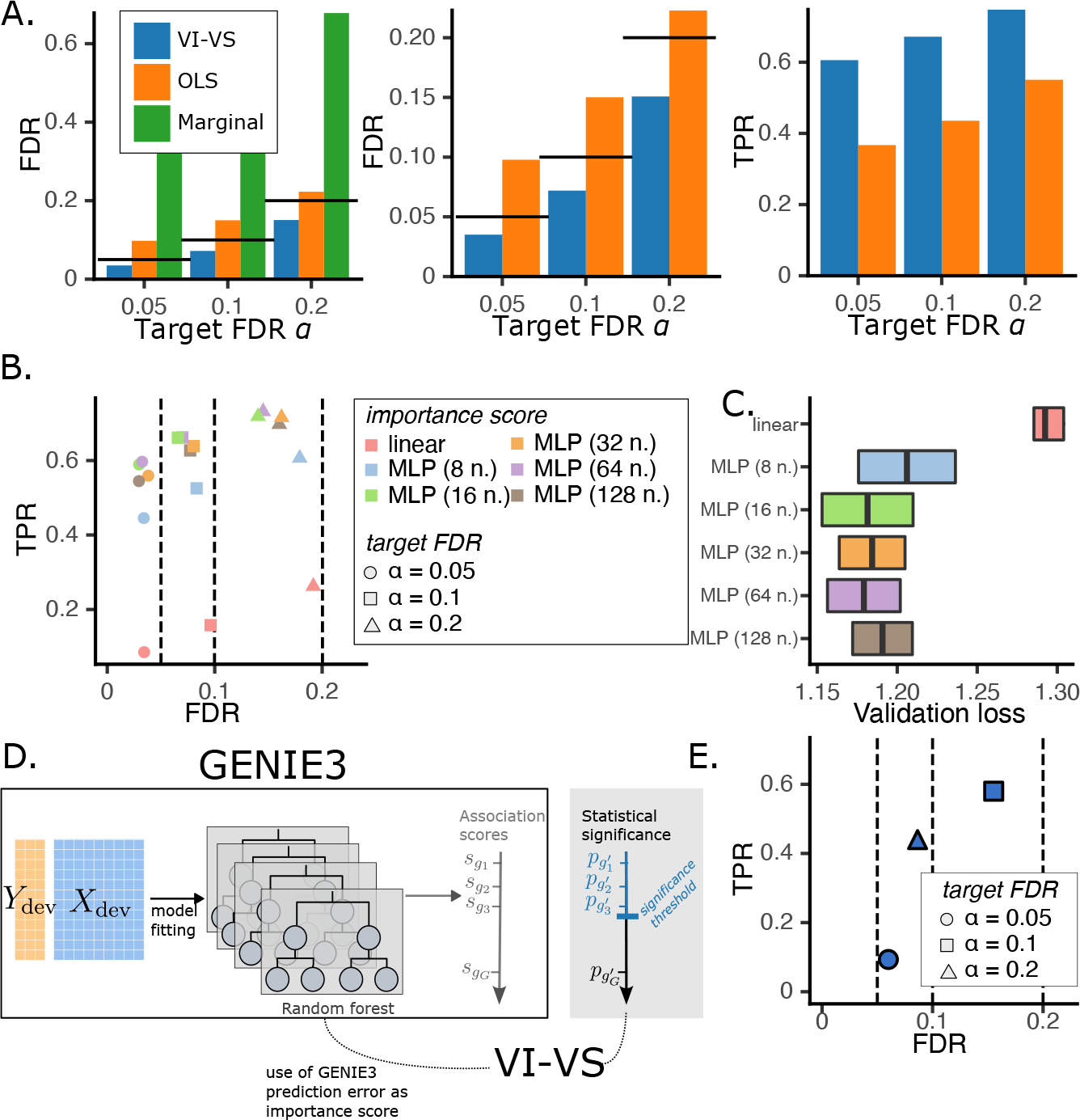
Semi-synthetic experiment. **A**. Comparison of FDR control and power for conditional independence testing at the *gene* level, averaged over five random weights initializations for the models of `VI-VS`, and across the five surface proteins of the dataset. Here, `VI-VS` uses a neural network with 64 units to compute feature importance scores. *Left*: FDR control comparison for the CRT, Ordinary Least Squares (OLS) under t-tests, and marginal independence tests. Because the marginal test did not control the FDR, it was removed from the rest of the experiments. *Center*: Zoom on the previous figure. *Right*: Associated TPR. **B**. FDR-TPR curves for different importance scores averaged over five random weights initializations for the models of `VI-VS`, and across the surface protein of the dataset. Circles, squares, and rectangles respectively represent the models’ decisions for target FDR levels of 0.05, 0.1, and 0.2. **C**. Associated held-out mean squared error of the different regression models used as importance scores. **D**. Use of `VI-VS` as a calibration tool for GENIE3. After fitting the regression tree ensemble of GENIE3, we used their prediction error as feature importance scores for `VI-VS`, allowing one to detect conditionally dependent genes with statistical significance. **E**. FDR/TPR levels of `VI-VS` using GENIE3 reconstruction losses as feature statistics. In this experiment only, for scalability reasons, we considered a total of 100 genes in the experiment. In **B**. and **E**., dashed lines denote target FDR levels.

The OLS performed better but still overestimated the FDR, possibly due to the violated linearity assumptions. Likelihood misspecification, i.e., invalid Gaussian assumptions on the data, is another risk that can cause FDR miscalibration for OLS. As an illustration, we repeated the simulation, this time generating the response variable counts from a Poisson distribution (Supplement D.1), in which case OLS performs worse. On the other hand, the application of `VI-VS` with an MLP for the importance score controlled the FDR in both these scenarios. We also evaluated the robustness of our approach to key characteristics of the simulated data, including sparsity and the number of genes in the assay, and show that our approach compares favorably to OLS, and provides consistent FDR control across all parameter settings (Supplement D.2).

#### Increased power using more complex importance score functions

Using better feature importance scores can increase the power of the CRT framework (i.e., lower type II error), while still maintaining calibration of the type I error estimates (as stated by the theoretical analysis in proposition 1 and [29]). To illustrate this, we considered a range of increasingly complex model choices to compute the importance scores and repeated our simulation analysis (Figure 4B). We found that all importance scores controlled the FDR at different levels. For instance, using linear regression with an OLS objective still controlled the FDR, although this model is a poor predictor of the response variable. Conversely, higher-capacity models learned with increasingly complex MLP architectures, led to increased power, indicating that more pertinent importance scores may lead to more discoveries. Selecting the most relevant importance score for a given application is not always clear, but we found that the models with the best held-out predictive performance also detected more true positives (Figure 4C). Therefore, we advise using predictive performance as a criterion to select the importance score to use with `VI-VS`.

#### Using VI-VS to calibrate existing algorithms for the identification of feature interactions in single-cell genomics

Our framework can build interpretable decisions on top of existing algorithms that lack a scalable or otherwise principled way to define calibrated decision rules. An example of such a model is GENIE3, which uses an ensemble of regression trees to produce scores that quantify the importance of each gene in predicting a held-out “response” gene, thus identifying putative interactions between genes. These scores were shown to have state-of-art performance in ranking putative interactions from the most to the least relevant. They, unfortunately, do not easily inform which interactions should be considered relevant for decision-making. Consequently, we used `VI-VS` to construct interpretable decisions on top of GENIE3. To do so, it suffices to plug in the GENIE3 model as our importance score in algorithm 1. Specifically, given a response variable *y*, we trained GENIE3 regression trees once using the development part of the data. We then used the respective prediction errors on the validation data as importance scores for `VI-VS` (Figure 4D; see Supplement B.4 for details). Application to our simulated data demonstrates that this wrapper procedure provides decision rules that control the FDR at several levels (Figure 4E), while still providing ample true positive rates. The CRT framework can therefore be used to better utilize a large family of algorithms in this area, as long as they produce an estimate of interaction “strength” that considers all features in *X* and can be used as a plugin importance function.

#### Multi-resolution testing as a way to increase power

The limited true positive rate in our simulation results can be explained to some extent by gene correlations that could not be resolved because of the limited size of the data. We next tested how accounting for different levels of resolution helps mitigate this. To this end, we applied our hierarchical procedure with different gene cluster granularities (here, 200 clusters, 300 clusters, or a single gene analysis; Figure 5A). We generally observed that the detections were consistent across the different resolutions, i.e., if a group of genes was identified at a given resolution, groups containing these genes were detected at coarser resolutions. Testing at multiple resolutions is useful to identify clusters of genes that were not detected at the gene level due to sample size limitations. Clusters to 6 are such examples, illustrating cases where relevant genes were not detected at the gene level, but were detected at coarser resolutions by `VI-VS`.

**Figure 5:**
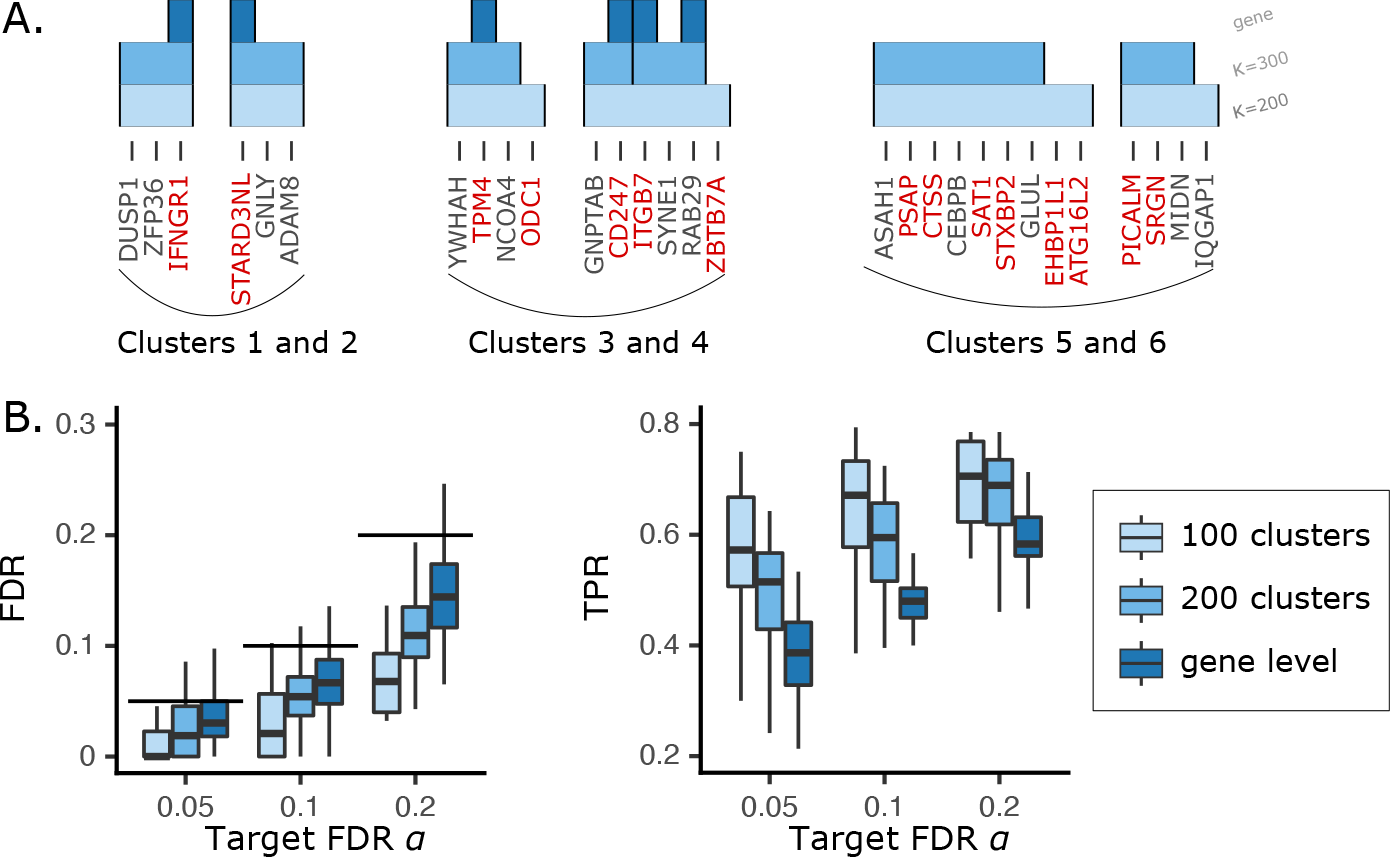
**A**. Examples of gene groups identified by `VI-VS` in the semi-synthetic experiment. Clusters 1 and 2 show examples where all genes affecting the surface protein expression in the simulation are detected at the gene level. Clusters 3 and 4 show examples where some of the genes are not detected at the gene level but are detected at coarser resolutions. Clusters 5 and 6 show examples where none of the genes are detected at the gene level, but are detected at coarser resolutions. **B**. FDR (*left*) and power (*right*) for `VI-VS` applied at different resolutions. When testing at the group level, a group of genes was considered a true positive if it contained at least one gene that was a true positive at the gene level.

For a more quantitative evaluation of the merits of multi-resolution testing, we computed FDR and power, where a selected group of genes was considered true positive if it contained at least one gene that was a true positive (at the gene level) and false positive otherwise (Figure 5B). Our results show that testing at coarser resolutions, i.e., grouping genes together, yielded more discoveries while maintaining calibrated FDR. More generally, multi-resolution testing provides more insight into the statistical relationships between genes and responses. When a coarse gene group is detected, the identification of finer-grained gene groups can be used to identify the individual genes that are responsible for the association [37]. Tying individual genes with coarser gene groups can also help identify and annotate genes with additional sources of information [38].

### 3.3 VI-VS identifies links between surface proteins and gene expression programs with CITE-seq

Next, we considered a CITE-seq dataset of PBMCs, obtained from eight healthy human donors [39]. We applied `VI-VS` at the single gene level, as well as the OLS baseline to a subset of this data with a total of 50,000 cells, 2,000 genes, and 224 surface proteins (Figure 6A). Notably, this dataset includes information from 13 batches, which could be accommodated by `VI-VS` due to the batch correction capacity built into its generative model [30]. Application of OLS to this case study (using an FDR cutoff of 10%) returned a very large number of associations for all 224 proteins (Figure S5A). In contrast, `VI-VS` (with the same target FDR) only predicted associations for 51 of the proteins, with a smaller number of interactions per protein (52 interactions on average, compared to 140 interactions with OLS). To understand this, we first considered the set of proteins that had no associations with `VI-VS`. Using TotalVI [40], we estimated for each protein the percentage of cells that plausibly express it on their surface (accounting for background signal, which is often observed in protein quantification with CITE-seq). We found that those proteins with no detection by `VI-VS` tend to have a much lower signal, compared to the ones that have been associated with gene expression (Figure S5B). Conversely, the OLS analysis identified associations for proteins that are likely not well captured or not expressed in these settings. OLS detected numerous associations (21, 41, 121, and 123 gene-protein pairs respectively) for four negative control proteins (Rat-IgG1-1, Rat-IgG2b, Rat-IgG1-2, and Rat-IgG2c) that are not expressed by human cells. In contrast, `VI-VS` detected none of these associations.

**Figure 6:**
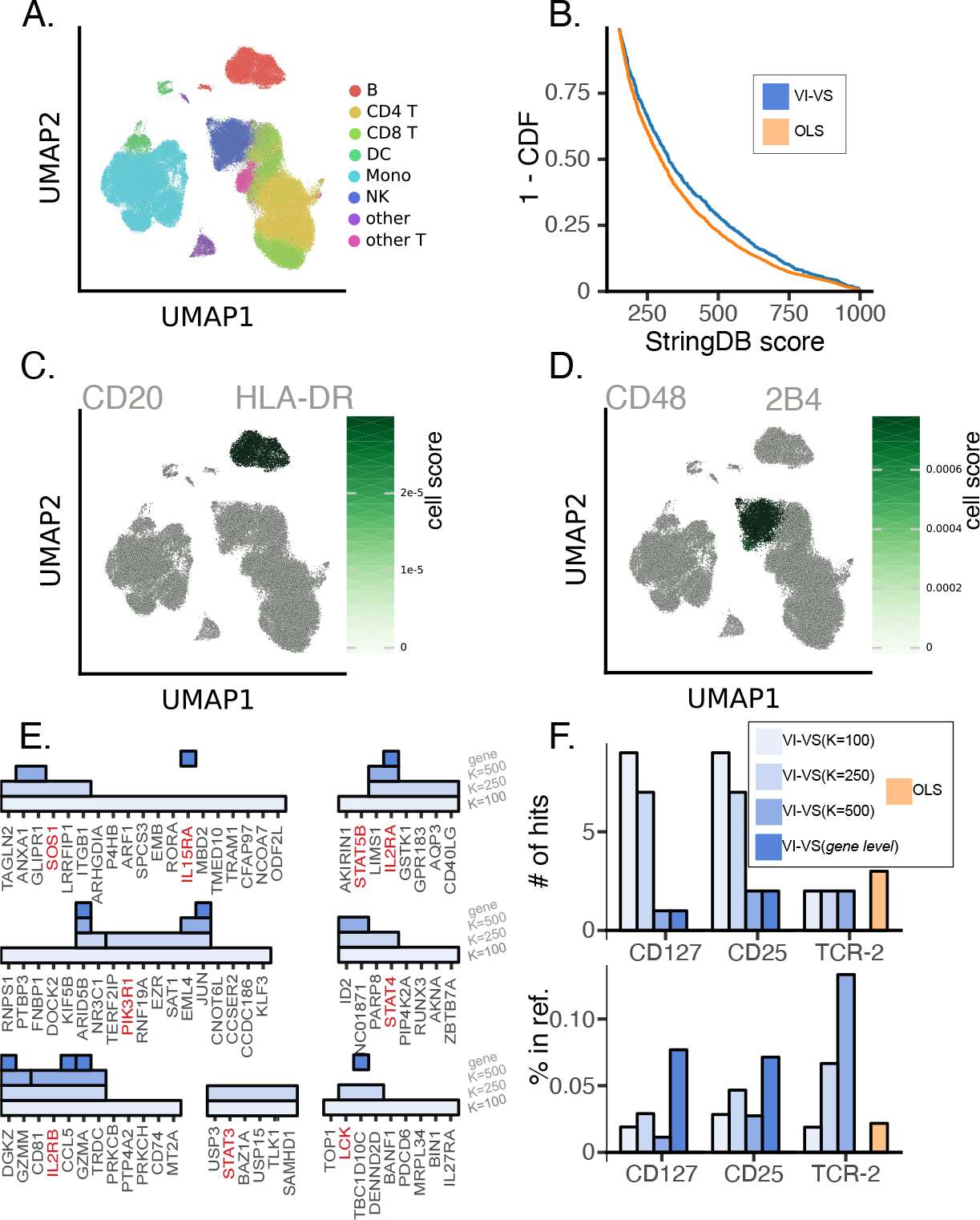
CiteSeq experiment. **A**. UMAP of the dataset. **B**. Distribution of stringDB scores of gene-protein discoveries made by `VI-VS` and least-squares (higher is better). **C**. and **D**. cell scores (averaged per cell-type) for the CD86-HLA and CD48-2b4 gene-protein pairs detected by `VI-VS`. High scores identify cells where the dependency is most expressed. **E**. Visualization of `VI-VS` detections at several resolutions for surface protein CD25 and T cells. Each filled rectangle characterizes a gene group detected as significant by `VI-VS` when testing for conditional independence at several resolutions (*K* ∈ *{*100, 250, 500) and at gene level). Genes in red correspond to genes contained in *Interleukin-2 Family Signaling R-HSA-451927* or *Interleukin-2 Signaling R-HSA-9020558* pathways. **F**. Agreement of the predictions with the Reactome pathway database, focusing on T cells for three surface proteins. *Top*: Number of predicted genes contained in each pathway, and *bottom*: proportion of predicted genes contained in each pathway over the total number of detections. The following pathways were considered: *Interleukin-7 Signaling R-HSA-1266695* for CD127, *Interleukin-2 Family Signaling R-HSA-451927* and *Interleukin-2 Signaling R-HSA-9020558* for CD25, and *TCR Signaling R-HSA-202403* for TCR-2. For **E**. and **F**., models were fit on T cells only.

To further compare the validity of the associations made by `VI-VS` and OLS, we used StringDB [41] to evaluate the a priori support for each interaction. Specifically, we assigned StringDB’s protein-protein “combined score” to each protein-gene pair. This combined score is a composite measure that integrates the scores of protein-protein associations computed across several modalities. We first compared the distribution of these scores for predicted gene-protein interactions across all proteins (Figure 6B). We found that the scores of the interactions predicted by `VI-VS` were significantly higher than the ones identified by OLS (Kolmogorov–Smirnov test, *P* ≤ 10^−6^). A similar trend was observed when comparing these scores for proteins for which both methods made predictions (Kolmogorov-Smirnov test, *P*≤ 0.05). This, combined with the fact that OLS detected associations for several negative control proteins, suggests that OLS is likely misspecified and may return many false positives. Conversely, `VI-VS` provides a more conservative and accurate way to identify biologically meaningful associations.

#### Locating protein-gene associations to the relevant cell subsets

A core feature of `VI-VS` is the ability to not only identify the association of genes with the response variable but also highlight the set of cells in which this interaction is more likely to be relevant. As a first example of this, we consider an association detected between MS41 (encoding the B cell marker CD20) and the HLA-DR receptor. Using the cell-specific importance scores from Equation 7, we identified B cells as the most relevant cells for this association (Figure 6C). This agrees with previous findings on the physical and functional association between CD20 and MHC-II in activated B cells [42], and the use of these two molecules as joint targets for combination therapy in lymphomas [43]. `VI-VS` also identified an association between the presentation of CD48 on the cell membrane and the expression of 2B4, which encodes the activating NK cell receptor CD244. The cell-specific importance scores suggest that this dependency is primarily driven by natural killer (NK) cells (Figure 6D). This result agrees with reports on the functional association between CD48 and CD244 in NK cells, where direct binding of these molecules is important to drive the surface expression and phosphorylation of CD244 in NK cells, consequently affecting their effector function [44].

When the practitioner has prior knowledge about the cell types of interest for the analysis, it is advantageous to fit the model only on these specific cells rather than the entire dataset. This choice reduces the computational cost of the algorithm and yields clean type-specific associations, eliminating the need for post-processing based on cell-specific importance scores. To illustrate how such `VI-VS` can unveil biologically relevant associations at multiple resolutions, we searched for associations between genes and proteins in a specific cell type specified before testing: T cells. We focused on the CD25 surface protein (IL2RA) and identified conditionally dependent genes and gene groups at different resolutions. For a coarse resolution (*K* = 100), `VI-VS` detected 26 groups of genes, seven of which contained genes known to be involved in the regulation of IL2RA [45], which we visualized in Figure 6E. In addition to IL2RA and IL2RB, detections at the gene level included CCR5, which encodes a chemokine receptor influencing IL2 production in T cells [46]. Testing at several resolutions simultaneously identified causal genes that were not detected at the gene level, presumably due to sample size limitations. For instance, STAT3 and STAT5B, two transcription factors involved in the regulation of IL2RA, were not detected by `VI-VS` at the gene level but detected at a coarser resolution. STAT3 promotes T cell survival and is known to inhibit T cell proliferation and IL2 production [47]. The activation of STAT5 by IL2 cytokines is a critical signaling pathway associated with regulatory T cell differentiation and function [48]. We generalized this analysis to other proteins, and compared the number of detected genes contained in known pathways for `VI-VS` and OLS more quantitatively (Figure 6F). `VI-VS` detections at coarser resolutions detected more overlapping genes contained in the pathways, while tests at finer resolutions provided more precise gene-level associations, and overall, more overlapping genes than OLS.

An important observation is that contrary to marginal approaches, `VI-VS` automatically controls for cell-type variation. A gene and a protein may be marginally dependent if they are expressed by the same cell types, even if they do not correlate within these types. Figure S6 compares, for every gene, significance scores for its association with the surface protein CD4, using a marginal test and `VI-VS` with a significance score of DE between CD4+ T cells and the rest of the cells. The marginal approach has a very strong correlation with the cell-type variation, while `VI-VS` does not. Consequently, a marginal test for gene-protein association may reflect cell-type variation rather than molecular interactions. Conversely, since gene expression data from all genes except one are sufficient to identify cell types, `VI-VS` conditions the variation between cell types and finds associations that are not explained by cell-type variation.

### 3.4 VI-VS identifies spatially-dependent gene expression programs in lymphocytes using ST

We showcased how `VI-VS` can be applied for spatial transcriptomic (ST) analyses. In particular, we studied an ST dataset consisting of one lung biopsy from a non-small cell lung cancer (NSCLC) patient, containing 960 genes and 200,000 cells and sequenced using the CosMx platform [49]. In this case, our objective was to link gene expression to spatial contexts that reflect cell localization in the tissue or its proximity to other cells.

#### Characterizing spatial differential expression patterns for T cells

We first aimed to identify differences in gene signature between T cells located in the tumor and lymphoid aggregates (Figure 7A). We trained the considered models on these cells using a binary response variable indicating the cell location (*y* = 1 for tumor cells, 0 for lymphoid aggregate cells); the importance score used by `VI-VS` was modified to the log-likelihood of a neural network binary classification model. We first compared the number of genes detected predicted by `VI-VS` to simple parametric and nonparametric differential expression tests (Table S1). The latter approaches detected almost all genes in the dataset. As the number of cells increases, negligible differences in gene expression are likely to be detected as significant, even after multiplicity correction. This is a known problem for point null hypothesis tests applied to single-cells [50, 51], which would require further filtering of the results to obtain a reasonable number of discoveries that can be interpreted. In contrast, `VI-VS` detected a much smaller number of genes at the gene level. Indeed, only five genes were detected by `VI-VS` at the gene level, including ITGAE and IL7R (Figure 7B). ITGAE, encoding CD103, is a canonical marker of tissue-resident memory CD8+ T cells (Trm). Its expression characterizes T cell infiltration in the tumor microenvironment (TME) [52]. Therefore, this result highlights the preferential state as resident memory T cells if they infiltrated the tumor. IL7R is a generic marker of CD4 memory T cells and is widely expressed in those cells. The multiresolution approach detected a larger set of genes known to capture known tumor-specific T cell signatures. ITGAE is associated with a group of genes that have a cytotoxic function in CD8 cytotoxic T cells (CTSW, GNLY, NKG7, PRF1, GZMB, KLRK1) up-regulated in the tumor. Studying this gene module therefore identifies resident CD8 Tcells in the tumor to have a highly cytotoxic and activated phenotype. IL7R on the contrary shows up in a module with CCR7 and KLF2, genes that mark naive CD4 Tcells. This module therefore identifies CD4 T cells to be enriched outside of the tumor. These genes were mainly located in spatial clusters of lymphocytes which we identified as lymphoid aggregates. In general, the genes and modules detected reflect a diverse set of biological processes varying across lymphoid aggregates and tumor regions.

**Figure 7:**
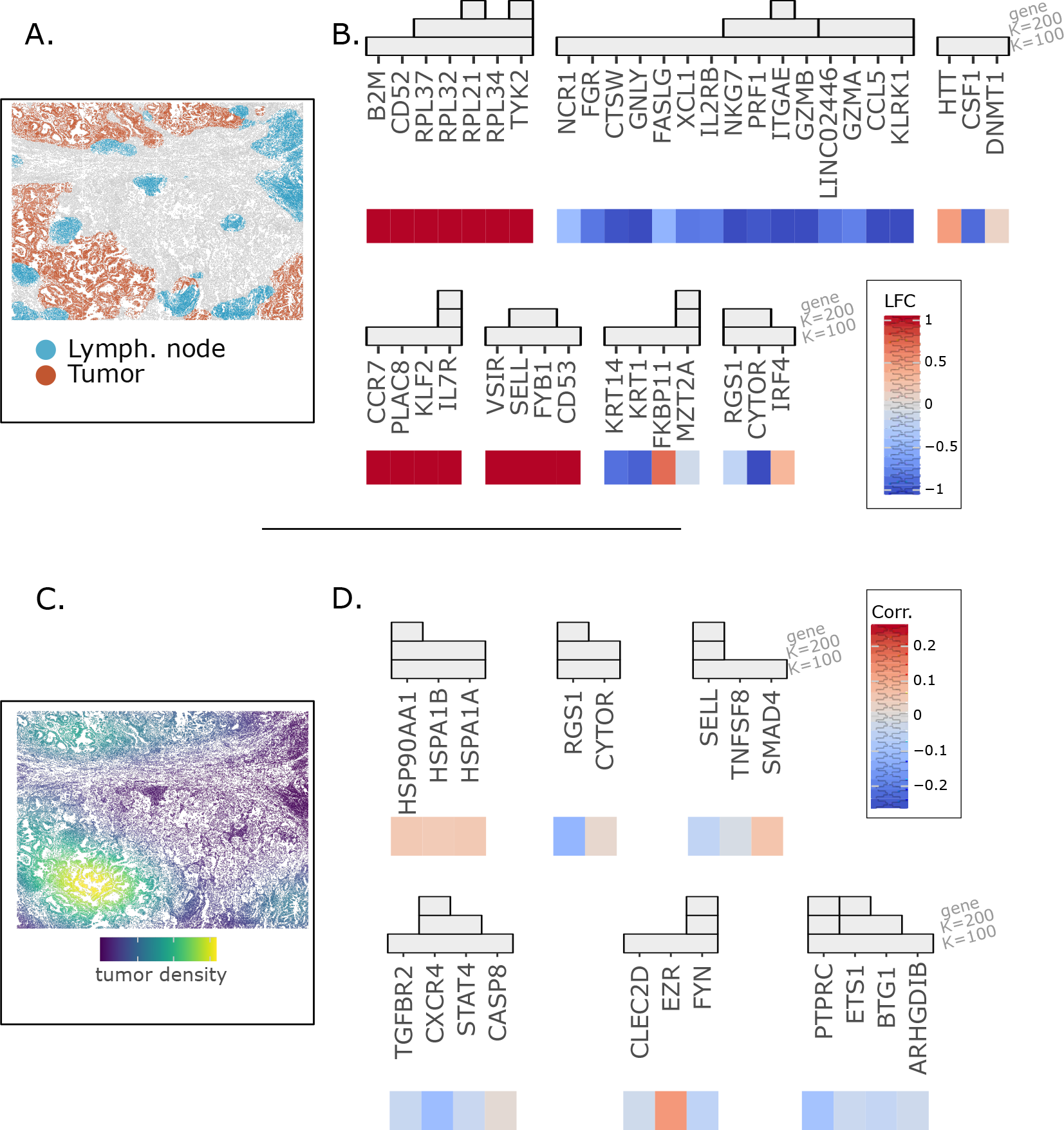
ST T cells experiment. **A**. Tissue segmentation into lymph nodes and tumor regions. **B**. Identified spatial DE genes by `VI-VS` in T cells, along with spatial LFCs (positive values denote gene upregulation in lymph nodes compared to tumors) against the significance scores of `VI-VS`. **C**. Local density of tumor cells in the tissue. Density is estimated using kernel density estimation (bandwidth of 500*μm*). **D**. Identified T cell genes conditionally associated with local density of tumor cells, along with marginal Spearman scores between gene expression and local tumor density.

#### Identifying T cell genes associated with tumor proximity

Last, we characterized the associations between gene expression and tumor proximity by defining *y* as the local density of tumor cells surrounding T cells. The first step was to define the range of tumor proximity we considered relevant. To do so, we constructed several responses *y*, corresponding to the predicted tumor density for each T cell predicted by a Gaussian kernel density estimator of different bandwidths, taking values in *h*∈ [10*μm*, 50*μm*, 100*μm*, 200*μm*, 500*μm*, 1000*μm*]. Of these, only *h* = 500*μm* and *h* = 1000*μm* detected associations at the gene resolution, suggesting that gene expression interactions with immediate and close tumor cells are more difficult to detect and would require more data to reach significance. We focused on the discoveries made by `VI-VS` for *h* = 500*μm* (Figure 7C), and visualized the detections, corresponding to six gene groups (Figure 7D). Our approach provided a significant number of genes related to T cell function in the tumor microenvironment, including HSP90AA1, ETS1, CXCR4, RGS1, and FYN, all detected at the gene level. HSP90AA1, for instance, encodes a heat shock protein, whose overexpression correlates with tumor progression and a poor prognosis in NSCLC [53, 54], and has been shown to correlate with an exhausted phenotype of CD8 T cells in tumors [55]. CXCR4 encodes a chemokine receptor whose expression is associated with the formation of lymphoid follicles that we detected outside of the tumor [56]. TGFBR2 is associated with this gene at the module level and has been shown to induce the residency of T cells in lymphoid tissue [57]. A correlation of RGS1 with T cell exhaustion has been observed in various cancers, including NSCLC [58]. We emphasize here that the location of T cells outside of the tumor is related to specific chemokine signals, specific cell states, and markers of exhaustion, whereas the location inside the tumor is related to an increase in heat shock protein signatures associated with T cell exhaustion and cellular stress. These results suggest that `VI-VS` is flexible to be applied to continuous descriptions of spatial localization and then helps to dissect function without prior knowledge of important tissue niches.

## 4 Discussion

`VI-VS` is a comprehensive framework for identifying potential functional relationships among molecular species in single-cell multiomics. It employs a nonparametric test for conditional independence, a concept that provides a more stringent notion of association than marginal tests. Unlike parametric tests, which require to posit a predefined relationship between features and the response, `VI-VS` does not require this relationship to be known. This makes `VI-VS` a versatile tool that remains valid even when the relationship between features and the response is unknown. `VI-VS` can be employed as a meta-algorithm to make the discoveries of existing methods more interpretable by constructing importance scores from their predictions. In this work, we calibrated GENIE3 discoveries via `VI-VS` but other models could be used instead. Used as a meta-algorithm, `VI-VS` could leverage the strengths of different models depending on the problem at hand.

Detecting conditional dependencies requires more data than identifying marginal dependencies [59]. This requirement may cause conditional approaches to miss potentially relevant associations due to limited statistical power. To address this risk, we proposed a multiresolution testing procedure. This procedure not only identifies individual genes with conditionally dependent features, but also recognizes feature groups that may contain them, providing a comprehensive characterization of the statistical dependencies between features and responses. We highlight throughout the manuscript that these modules can help in identifying the functional role of an identified molecule and thereby help in interpreting the results.

In the general case, `VI-VS` discoveries have no guarantees to be functional. Feedback loops prevalent in molecular interactions, cell communication, or unobserved molecular species are just a few examples of phenomena that lead to spurious discoveries. However, certain multiomic setups already offer promising avenues for identifying causal relationships with `VI-VS`. Identifying reproducible and robust discoveries across biogically diverse environments could help mitigate the effect of unobserved confounders [60, 61]. Furthermore, causal candidates identified by `VI-VS` could be validated through interventional experiments such as Perturb-seq [62]. Making no assumption on the distribution of the response, our approach can readily be applied to other multiomic setups. A potential application could be the identification of gene associations with metabolites [63]. `VI-VS` could also be applied more broadly to spatial transcriptomics to create complex characterizations of cell phenotypes and their environments. It could, for instance, pinpoint genes involved in receptor-ligand interactions [64], or in determining cellular morphologies [65]

The presence of technical data variation, including differences in sample preparation or sequencing technologies, is a major challenge for large-scale multiomic analysis [66, 67]. `VI-VS` effectively addresses this issue by conditioning on these nuisance factors. As the generative models can capture multiple nonlinear technical effects [68], our approach can produce robust discoveries in complex settings [69]. Therefore, we propose `VI-VS` as a general framework to yield estimates of effect sizes in those complex settings, and `VI-VS` thereby produces robust findings by consolidating large datasets across multiple batches [69].

## Code availability

The code to reproduce the results in this paper is available at

https://github.com/PierreBoyeau/VIVS-reproducibility.

## Acknowledgments and Disclosure of Funding

N.Y. is an advisor and/or has equity in Cellarity, Celsius Therapeutics, and Rheos Medicine.

## Supplement

### A Theoretical background and guarantees

#### A.1 False Discovery Rate control

FDR control aims at ensuring that the rate of false positive detections made in multiple hypothesis testing scenarios is controlled in expectation. Formally, the FDR characterizes the expected proportion of erroneous discoveries:

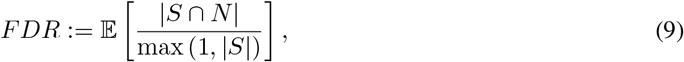

where *S* denotes the set of detected associations, and *N* the set of non-existing associations in the data.

#### A.2 Proof of Proposition 1

Proposition 1 is a consequence of the following theorem,

##### Theorem 1

(CRT gives valid inference, from Lemma 4.1 in [29]). *Let* 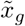 *denote some synthetic sample for feature g from some proposal distribution (a distribution used to generate synthetic samples). Suppose under* 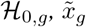 *satisfy the following property:*

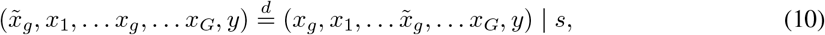

*for k* = 1, …, *K. Above, the notation* 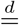 *denotes equality in distribution conditioned to s. If* 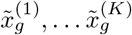 *are K i*.*i*.*d. draws from this proposal distribution, then*,

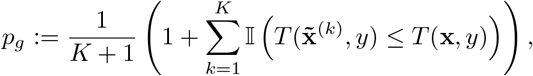

*where* 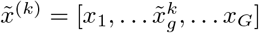, *k* ≤ *K, is a valid p-value for* ℋ_0,*g*_ *(i*.*e*., *for the conditional independence test)*.

Property (10) ensures that the distribution of the data remains the same even when the true and synthetic samples are swapped. Equation 10 is a pairwise exchangeability property that ordered sets 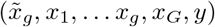 and its swapped counterpart 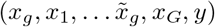 have the same distribution. In particular, this property is a relaxation of the exchangeability propert required to construct knockoffs. We refer to [29] for a more detailed discussion and interpretation of this property.

*Proof*. Assume that the distribution of synthetic samples 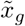 satisfy Property (10) under the null. Then, conditional on *s*, under the null we have

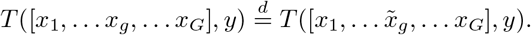

Since 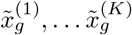 are i.i.d draws from the conditional distribution and *x*_g_ is an independent sample from the same conditional distribution, we conclude

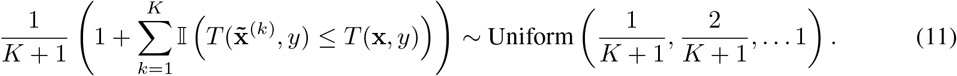

□

We now use this result to prove Proposition 1, and in particular, that the two described sampling schemes verify the exchangeability property required to apply Theorem 1.

*Proof*. Proposition 1 states that two sampling schemes are valid for the CRT. The first sampling scheme corresponds to sampling from the distribution given by ∫*p*(*x*_*g*_| *z, s*)*p*(*z* |*x*_− *g*_, *s*)*dz*. From our assumptions on the generative model, we have that

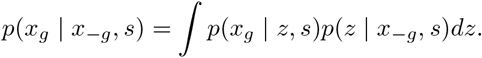

Hence, (10) is satisfied. (See [29] for a more detailed proof.)

We now consider the second sampling scheme, which is the case presented in Algorithm 1. We let 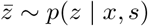 be a latent variable sampled from the posterior distribution. Let *x* := [*x*_1_, … *x*_*g*_, … *x*_*G*_] denote the observed gene expression profile. For clarity in the following equations, the subscript notation is used to specify which random variable is being considered. For example, 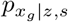 represents the density of random variable *x*_*g*_ given *z* and *s*.

Under the null, we have the following joint distribution

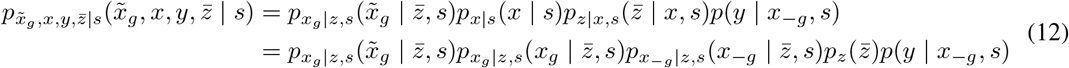

These equalities follow from the assumed factorization of the generative model and from Bayes’ rule. Notice the last expression is symmetric in the arguments *x*_*g*_ and 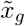 . We conclude that *x*_*g*_ and 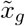 have the same distribution conditional on *s* and 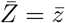 . We can thus apply Theorem 1 with conditioning on 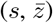. □

Observed that exchangeability property does not hold unconditional to a fixed value of *z*. This can be seen in Equation 12, since 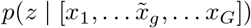 and *p*(*z* | [*x*_1_, … *x*_*g*_, … *x*_*G*_]) are not equal in general. This justifies why *z* must be fixed when sampling 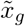, and cannot be sampled from the posterior distribution at every MC trial.

### B Supplementary Information on VI-VS

#### B.1 Multiresolution analysis

In this section, we describe how `VI-VS` performs multiresolution analysis.

##### Feature clustering

The first step of this procedure consists in clustering features. While any clustering procedure can be used, we rely on a custom hierarchical clustering procedure when features are genes. After fitting scVI on all genes, we compute cell and gene specific denoised gene expresssion levels from the fitted model. We then compute the empirical correlation matrix between genes. We then used complete-linkage clustering to cluster genes at arbitrary resolution.

##### Multiresolution testing

Testing for conditional independence at the cluster level follows the same procedure as in the main text. In particular, let *A* denotes a set of genes for which we want to test

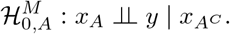

We first randomize the expression for all genes in *A*, obtaining expressions 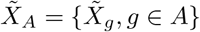, where 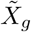 is obtained from Equation 6. We then construct the randomized expression profile for all genes,

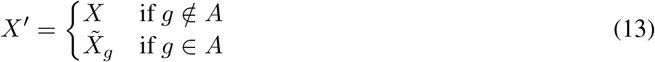

We then compute the randomized test statistic as *T* (*X*^*′*^, *Y, S*). We then repeat this procedure *K* times, obtaining *K* randomized test statistics *T* ^(1)^, … *T* ^(*K*)^, which we compare with the actual test statistic to construct the p-value for the conditional null.

##### Computational considerations

`VI-VS` relies on an efficient implementation that allows one to compute p-values across multiple resolutions in parallel. In particular, we use the same synthetic samples across all resolutions.

#### B.2 Implementation of VI-VS

We implemented `VI-VS` in python using Jax as backend. In particular, we relied on Algorithm S1 to compute the p-values, allowing to efficiently compute p-values using GPU acceleration. Our implementation relies on just-in-time compilation and parallelized computation to achieve high performance. This choice provides a two-fold improvement over an implementation relying on Pytorch, that benefits from a similar GPU acceleration but does not support just-in-time compilation nor parallelized computation.

##### Algorithm S1 Minibatched VI-VS

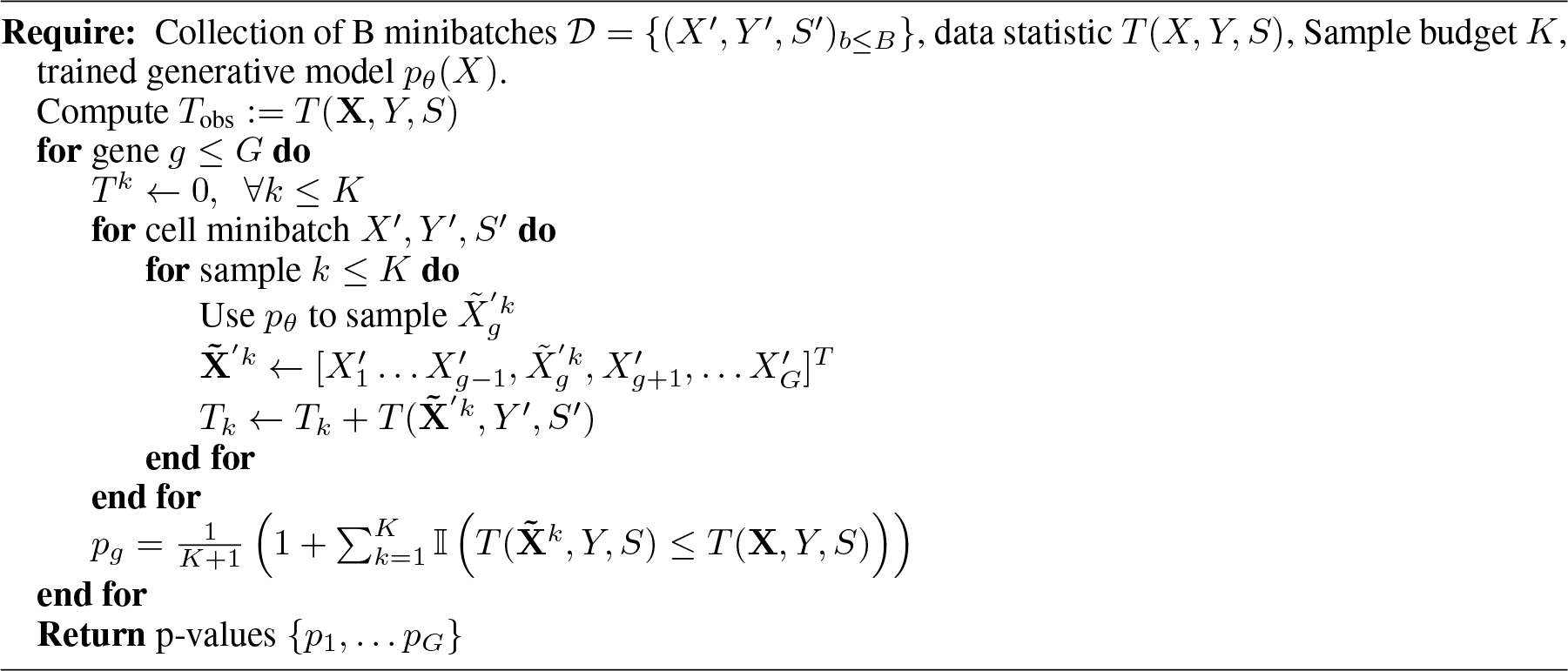

#### B.3 Generalization to multidimensional *Y*

While the main text of this work assumes that *Y* is unidimensional, we here outline how to handle cases where the measurement is *D*-dimensional. To do so, we consider a multivariate regression task to fit the importance score. We train its parameters via optimization of the data likelihood, which writes for a single observation as 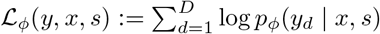, assuming conditional independence of the features of *y*.

Two scenarios can appear when performing conditional independence testing with `VI-VS` in a multidimensional setup. First, it can be desirable to retrieve conditional dependencies specific to each feature of *y*,

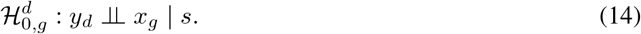

In this case, we use 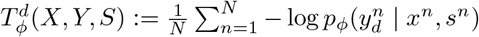 as importance scores to test 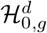 . It can also be the case that we would like to test

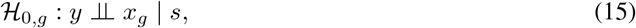

where in this case *y* is multidimensional. In this scenario, we use the full likelihood of the predictive model *p*_*ϕ*_ as importance scores.

#### B.4 Using VI-VS as a way to calibrate an existing feature selection method

`VI-VS` can be used to calibrate some existing feature selection methods that would not provide significance scores. Methods which rely on a regression fit to the data to ultimately score interactions between features and the response can straightforwardly be calibrated with `VI-VS`. Such methods include, for instance, regularized linear regression models for which significance scores are not easily available [21, 70], or ensembling methods [16, 17], that are popular feature selection tools employed for GRN inference.

For such methods, we first fit the regression model *f* on the development split, and define the importance score for the CRT as the mean prediction error of the model:

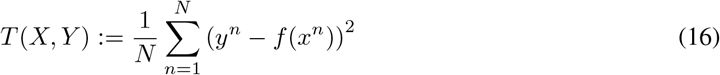

### C Supplementary Information on the experiments

#### C.1 Models

`VI-VS` We trained both scVI and the model associated with the importance scores until convergence using early stopping criteria [71] on the held-out validation data.

##### OLS and marginal tests

We computed OLS and marginal tests from log-CPM normalized counts. Tests relative to OLS and to the marginal baseline were fitted using the *statsmodels* python package [72].

##### Multiplicity control

P-values obtained with `VI-VS` OLS, and the marginal test were adjusted using the Benjamini-Hochberg procedure [33].

#### C.2 Semi-synthetic data

We consider a PBMC scRNA-seq dataset consisting of *N* = 6, 855 cells and *G* = 500 genes. We construct 5-dimensional synthetic surface protein measurements using the following scheme. Let *p* ≤ 5 be some protein. We first construct the subset of conditionally dependent genes *S* using Binomial sampling, with sparsity rate *s* = 0.3. Mean measurements for protein *p* are obtained as

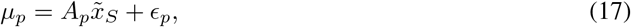

*A*_*p*_ ∈ ℝ^*N×s*^ corresponds to the ground-truth regressors, constructed as *A*_*p*_ = *βB*, where *B* denotes the *N* × *s* matrix of ones and *β* is the signal strength, parameterized as

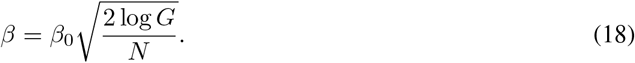

#### C.3 CITE-seq experiment

##### Data preprocessing

The original data [39] contained gene expression and surface protein counts for eight donors at several time points, corresponding to both pre-vaccination and post-vaccination. We focused our analysis on the pre-vaccination time points, resulting in a dataset containing a total of 50,000 cells and 2,000 genes. These genes were selected based on a procedure hoping to retain as much information about cells’ biological states as possible. To do so, we selected genes based on their joint variability with other genes instead of their marginal variability. To do so, we first clustered the genes using k-means based on their empirical correlation matrix, computed on log-CPM counts. In each cluster, we selected the gene with the highest highly variable-gene score, as predicted by Seurat [73]. The surface protein measurements *y* were centered and scaled to have zero mean and unit variance.

#### C.4 Nanostring experiment

##### Data segmentation and annotation

Baysor [74] was used to segment the tissue into individual cells. We downloaded the lung FFPE Nanostring CosMX data [75]. We used the position file of Lung 5 replicate 2 and split this into separate files for each FOV. Inside each FOV, we used the provided segmentation from Nanostring as a prior for Baysor segmentation and chose 0.1 as the prior segmentation weight. In short, we multiplied the z-position of each molecule by 10 to adjust for anisotropic distances. These results were concatenated across all FOV and scVI [30] was used to integrate these field of views. Leiden clustering was performed after neighbor calculation in latent space and cell-types were annotated based on known marker genes. We focused our analysis on T cells.

##### Identification of T cell markers in spatial analysis

A number of genes not expressed in T cells might still be detected in the T cell analysis, due to contamination events, e.g., due to imperfect cellular segmentation. To circumvent this issue, we restrict the analysis of the spatial datasets to genes that are higher expressed in T cells compared to other cell-types. We identified these genes via one-vs-all differential expression analysis based on log-CPM counts, using t-tests to compare mean expression levels, and BH to correct for multiple testing. We then constructed a list of 245 genes, containing all genes with adjusted p-values *<* 0.05, and with a log-fold change *>* 0.1 (upregulated in T cells).

### D Additional results

#### D.1 Semisynthetic Poisson experiment

We also considered a scenario where protein expressions suffer from Poisson instead of Gaussian noise, and where the relationship the protein expressions means are a linear combination of *squared* gene expressions. In such a scenario, an approach based on OLS does not produce valid p-values because of the model assumptions’ violations, while our approach still returns valid p-values (Figure S3). The power of our approach remains high, and has an higher area under the precision-recall curve to OLS.

#### D.2 Influence of key parameters on the semi-synthetic experiment

Contrary to OLS, the CRT-based approach provides consistently calibrated predictions (Figure S4). We however note that our approach displays intriguingly high FDR values for different sparsity levels. These scenarios correspond to regimes where the test detects a handful of discoveries, where we expect SeqStep to be unreliable.

#### D.3 Additional CITE-seq experiments

Figure S5 shows that the proteins for which `VI-VS` detects conditional associations have a higher percentage of cells expressing the protein.

#### D.4 Comparison of the number of significant hits for the spatial DE analysis

Table S1 compares the number of significant hits for all considered methods for the spatial DE analysis.

**Figure S1:**
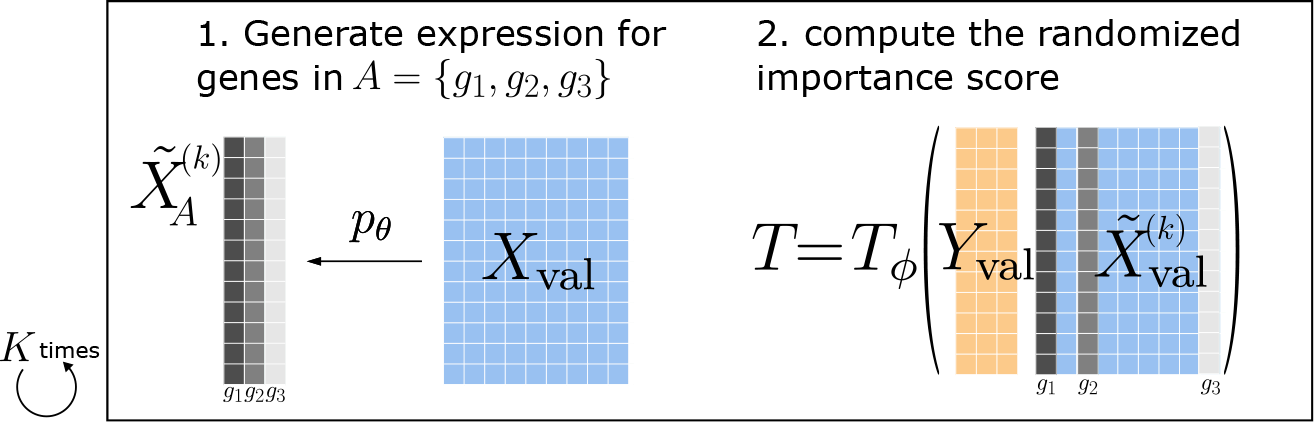
Illustration of the randomization procedure when testing at the gene group level. We rely on the generative model to produce synthetic gene expression profiles for all genes in the group *A*. Then, the randomized values are replaced in the original gene expression matrix to compute the randomized expression scores. This procedure is repeated *K* times to obtain the p-value for the conditional null.

**Figure S2:**
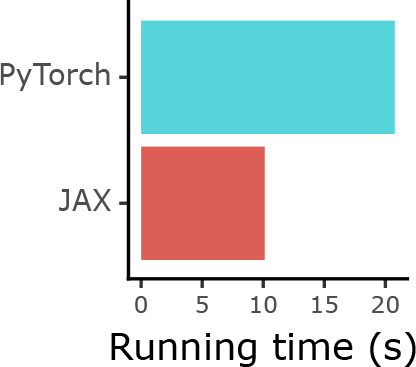
Running time comparisons. Execution time of `VI-VS` on a single GPU (RTX 3090) to compute p-values on a dataset containing 6,000 observations and 500 genes for 100 MC trials.

**Figure S3:**
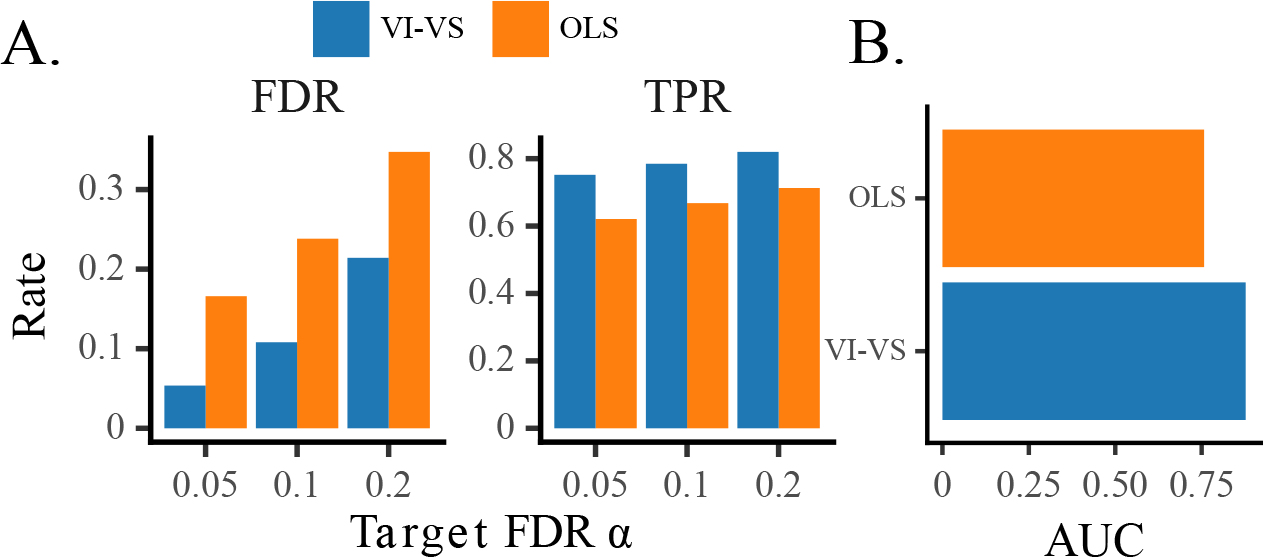
Semisynthetic experiment from a Poisson model. **A**. *Left*: FDR levels reached by `VI-VS` and OLS. *Right*: Power levels reached by `VI-VS` and OLS. **B**. Comparison of the area under the precision-recall curves of the different algorithms.

**Figure S4:**
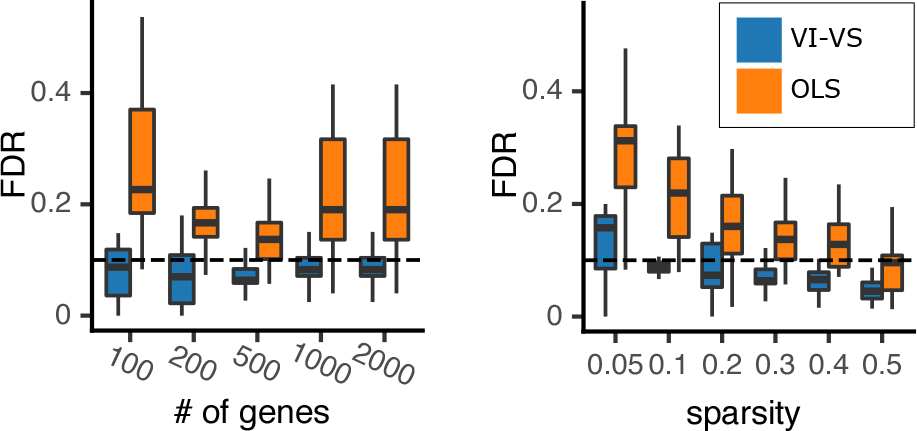
Influence of several parameters of importance in the semi-synthetic experiment. *Left*: Influence of the number of observed genes in the experiment. *Right*: Influence of the sparsity of the ground-truth parameter *β*.

**Figure S5:**
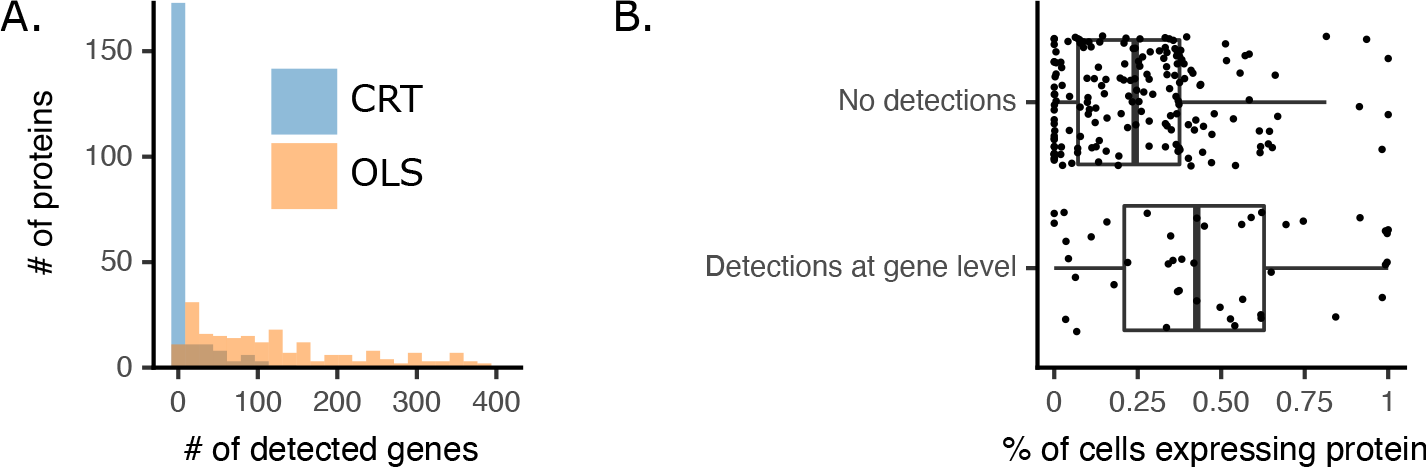
**A**. Comparison of the number of detections made by `VI-VS` and OLS across all proteins. **B**. Comparison of the percentage of cells expected to express the proteins for proteins for which (i). `VI-VS` predicts associated genes, and (ii). `VI-VS` does not predict associated genes. We predicted the percentage of cells expressing the proteins using TOTALVI, defining that a cell expressed a protein when the posterior probability observed protein counts comes from the foreground mode was above 0.95. Differences between the two groups are statistically significant (p-value *<* 10^−4^ under a Kolmogorov–Smirnov test). In both these experiments, a gene-protein association is considered significant when the adjusted p-value is below 0.1.

**Figure S6:**
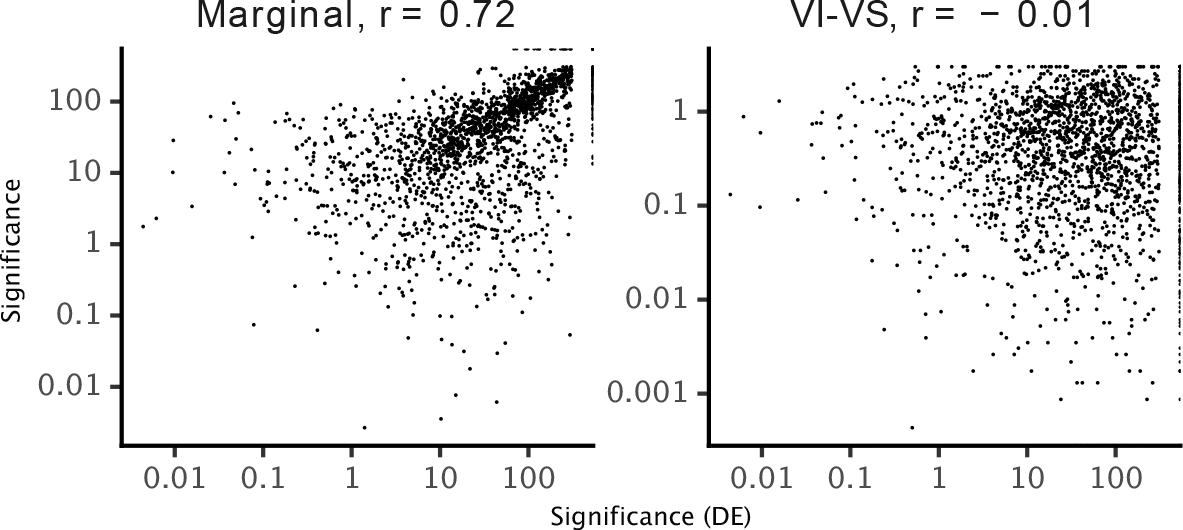
Comparison of significance scores for a Wilcoxon test between CD4+ and CD4-cells (*x-axis*) gene-protein significance scores (*y-axis*) for both a marginal test (*left*) and `VI-VS` (*right*). Significance scores were obtained as negative log p-values for the various tests. Correlation scores were computed using Spearman’s rank correlation.

**Table S1:**
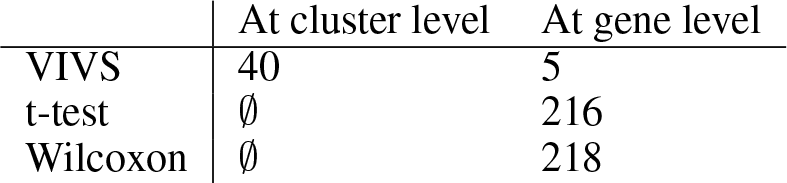
Comparison of the number of significant hits for the spatial DE analysis for `VI-VS` as well as common DE approaches.

## Footnotes

1 Here, superscripts in parentheses denote Monte Carlo samples.

2 https://github.com/YosefLab/VIVS

## References

1. Stoeckius, M. et al. Simultaneous epitope and transcriptome measurement in single cells. en. Nature Methods 14, 865–868 (Sept. 2017).

2. Lareau, C. A. et al. Droplet-based combinatorial indexing for massive-scale single-cell chromatin accessibility. en. Nature Biotechnology 37, 916–924 (Aug. 2019).

3. Chen, A. et al. Spatiotemporal transcriptomic atlas of mouse organogenesis using DNA nanoball-patterned arrays. en. Cell 185, 1777–1792.e21 (May 2022).

4. Wang, G., Moffitt, J. R. & Zhuang, X. Author Correction: Multiplexed imaging of high-density libraries of RNAs with MERFISH and expansion microscopy. en. Scientific Reports 8, 6487 (Apr. 2018).

5. Tornow, S. & Mewes, H. W. Functional modules by relating protein interaction networks and gene expression. en. Nucleic Acids Research 31, 6283–6289 (Nov. 2003).

6. Moses, L. & Pachter, L. Museum of spatial transcriptomics. Nature Methods, 1–13 (2022).

7. Segal, E. et al. Module networks: identifying regulatory modules and their condition-specific regulators from gene expression data. en. Nature Genetics 34, 166–176 (May 2003).

8. Iancu, O. D. et al. Utilizing RNA-Seq data for de novo coexpression network inference. Bioinformatics 28, 1592–1597 (June 2012).

9. Hu, R., Qiu, X., Glazko, G., Klebanov, L. & Yakovlev, A. Detecting intergene correlation changes in microarray analysis: a new approach to gene selection. BMC Bioinformatics 10, 1–9 (2009).

10. Van Dam, S., Võsa, U., van der Graaf, A., Franke, L. & de Magalhães, J. P. Gene co-expression analysis for functional classification and gene–disease predictions. en. Briefings in Bioinformatics 19, 575–592 (Jan. 2017).

11. Yang, L. et al. scMAGeCK links genotypes with multiple phenotypes in single-cell CRISPR screens. en. Genome Biology 21, 19 (Jan. 2020).

12. Causal Inference for Statistics, Social, and Biomedical Sciences (Cambridge University Press, 2015).

13. Gillis, J. & Pavlidis, P. “Guilt by association” is the exception rather than the rule in gene networks. PLoS Computational Biology 8, e1002444 (2012).

14. Margolin, A. A. et al. ARACNE: an algorithm for the reconstruction of gene regulatory networks in a mammalian cellular context. en. BMC Bioinformatics 7, S7 (Mar. 2006).

15. Lee, H. K., Hsu, A. K., Sajdak, J., Qin, J. & Pavlidis, P. Coexpression analysis of human genes across many microarray data sets. Genome Research 14, 1085–1094 (June 2004).

16. Huynh-Thu, V. A., Irrthum, A., Wehenkel, L. & Geurts, P. Inferring regulatory networks from expression data using tree-based methods. en. PLoS One 5 (Sept. 2010).

17. Moerman, T. et al. GRNBoost2 and Arboreto: efficient and scalable inference of gene regulatory networks. en. Bioinformatics 35, 2159–2161 (June 2019).

18. Kim, S. ppcor: An R Package for a Fast Calculation to Semi-partial Correlation Coefficients. en. Communications for Statistical Applications and Methods 22, 665–674 (Nov. 2015).

19. Chan, T. E., Stumpf, M. P. H. & Babtie, A. C. Gene Regulatory Network Inference from Single-Cell Data Using Multivariate Information Measures. en. Cell Systems 5, 251–267.e3 (Sept. 2017).

20. Qiu, X. et al. Inferring Causal Gene Regulatory Networks from Coupled Single-Cell Expression Dynamics Using Scribe. en. Cell Systems 10, 265–274.e11 (Mar. 2020).

21. Tibshirani, R. Regression shrinkage and selection via the lasso. en. Journal of the Royal Statistical Society 58, 267–288 (Jan. 1996).

22. Meinshausen, N. & Bühlmann, P. High-dimensional graphs and variable selection with the Lasso. The Annals of Statistics 34, 1436–1462 (June 2006).

23. Pratapa, A., Jalihal, A. P., Law, J. N., Bharadwaj, A. & Murali, T. M. Benchmarking algorithms for gene regulatory network inference from single-cell transcriptomic data. en. Nature Methods 17, 147–154 (Feb. 2020).

24. Peters, J., Janzing, D. & Schölkopf, B. Elements of causal inference: foundations and learning algorithms (The MIT Press, 2017).

25. Krishnaswamy, S. et al. Systems biology. Conditional density-based analysis of T cell signaling in single-cell data. en. Science 346, 1250689 (Nov. 2014).

26. Melenhorst, J. J. et al. Decade-long leukaemia remissions with persistence of CD4+ CAR T cells. Nature 602, 503–509 (2022).

27. Sacco, K. et al. Immunopathological signatures in multisystem inflammatory syndrome in children and pediatric COVID-19. Nature Medicine 28, 1050–1062 (2022).

28. Van de Sande, B. et al. A scalable SCENIC workflow for single-cell gene regulatory network analysis. Nature Protocols 15, 2247–2276 (2020).

29. Candes, E., Fan, Y., Janson, L. & Lv, J. Panning for gold:’model-X’knockoffs for high dimensional controlled variable selection. Journal of the Royal Statistical Society: Series B (Statistical Methodology) 80, 551–577 (2018).

30. Lopez, R., Regier, J., Cole, M. B., Jordan, M. I. & Yosef, N. Deep generative modeling for single-cell transcriptomics. Nature Methods 15, 1053–1058 (2018).

31. Lotfollahi, M., Wolf, F. A. & Theis, F. J. scGen predicts single-cell perturbation responses. en. Nature Methods 16, 715–721 (July 2019).

32. Ding, J. & Regev, A. Deep generative model embedding of single-cell RNA-Seq profiles on hyperspheres and hyperbolic spaces. en. Nature Communications 12, 2554 (May 2021).

33. Benjamini, Y. & Hochberg, Y. Controlling the False Discovery Rate: A Practical and Powerful Approach to Multiple Testing. The Journal of the Royal Statistical Society, Series B 57, 289–300 (1995).

34. Sesia, M., Katsevich, E., Bates, S., Candès, E. & Sabatti, C. Multi-resolution localization of causal variants across the genome. Nature Communications 11, 1093 (2020).

35. DeTomaso, D. & Yosef, N. Hotspot identifies informative gene modules across modalities of single-cell genomics. Cell Systems 12, 446–456 (2021).

36. Zheng, G. X. Y. et al. Massively parallel digital transcriptional profiling of single cells. en. Nature Communications 8, 14049 (Jan. 2017).

37. Schaid, D. J., Chen, W. & Larson, N. B. From genome-wide associations to candidate causal variants by statistical fine-mapping. Nature Reviews Genetics 19, 491–504 (2018).

38. Liberzon, A. et al. The molecular signatures database hallmark gene set collection. Cell Systems 1, 417–425 (2015).

39. Hao, Y. et al. Integrated analysis of multimodal single-cell data. Cell 184, 3573–3587 (2021).

40. Gayoso, A. et al. Joint probabilistic modeling of single-cell multi-omic data with totalVI. en. Nature Methods 18, 272–282 (Feb. 2021).

41. Franceschini, A. et al. STRINGdb package vignette. Nucleic Acids Research (2013).

42. Léveillé, C., AL-Daccak, R. & Mourad, W. CD20 is physically and functionally coupled to MHC class II and CD40 on human B cell lines. European Journal of Immunology 29, 65–74 (Jan. 1999).

43. Zeng, J., Liu, R., Wang, J. & Fang, Y. A bispecific antibody directly induces lymphoma cell death by simultaneously targeting CD20 and HLA-DR. Journal of Cancer Research and Clinical Oncology 141, 1899–1907 (Nov. 2015).

44. Claus, M., Wingert, S. & Watzl, C. Modulation of natural killer cell functions by interactions between 2B4 and CD48 in cis and in trans. Open Biology 6 (May 2016).

45. Jassal, B. et al. The reactome pathway knowledgebase. Nucleic Acids Research 48, D498–D503 (2020).

46. Camargo, J. F. et al. CCR5 expression levels influence NFAT translocation, IL-2 production, and subsequent signaling events during T lymphocyte activation. The Journal of Immunology 182, 171–182 (Jan. 2009).

47. Oh, H.-M. et al. STAT3 protein promotes T-cell survival and inhibits interleukin-2 production through up-regulation of Class O Forkhead transcription factors. Journal of Biological Chemistry 286, 30888–30897 (Sept. 2011).

48. Mahmud, S. A., Manlove, L. S. & Farrar, M. A. Interleukin-2 and STAT5 in regulatory T cell development and function. JAKSTAT 2, e23154 (Jan. 2013).

49. He, S. et al. High-plex multiomic analysis in FFPE tissue at single-cellular and subcellular resolution by spatial molecular imaging. bioRxiv [Preprint], 2021–11 (2021).

50. Love, M. I., Huber, W. & Anders, S. Moderated estimation of fold change and dispersion for RNA-seq data with DESeq2. en. Genome Biology 15, 550 (2014).

51. Boyeau, P. et al. An empirical Bayes method for differential expression analysis of single cells with deep generative models. Proceedings of the National Academy of Sciences 120, e2209124120 (2023).

52. Hu, X. et al. ITGAE Defines CD8+ Tumor-Infiltrating Lymphocytes Predicting a better Prognostic Survival in Colorectal Cancer. EBioMedicine 35, 178–188 (Sept. 2018).

53. Garcia-Carbonero, R., Carnero, A. & Paz-Ares, L. Inhibition of HSP90 molecular chaperones: moving into the clinic. The Lancet Oncology 14, e358–e369 (2013).

54. Esfahani, K. & Cohen, V. HSP90 as a novel molecular target in non-small-cell lung cancer. Lung Cancer: Targets and Therapy, 11–17 (2016).

55. Yuan, Z., Wang, L. & Chen, C. Analysis of the prognostic, diagnostic and immunological role of HSP90α in malignant tumors. Frontiers in Oncology 12, 963719 (2022).

56. Nagy, N. et al. In and out of the bursa—the role of CXCR4 in chicken B cell development. Frontiers in Immunology 11, 1468 (2020).

57. Li, G. et al. TGF-β-dependent lymphoid tissue residency of stem-like T cells limits response to tumor vaccine. Nature Communications 13, 6043 (2022).

58. Bai, Y., Hu, M., Chen, Z., Wei, J. & Du, H. Single-cell transcriptome analysis reveals RGS1 as a new marker and promoting factor for T-cell exhaustion in multiple cancers. Frontiers in Immunology, 5153 (2021).

59. Hastie, T., Tibshirani, R., Friedman, J. H. & Friedman, J. H. The elements of statistical learning: data mining, inference, and prediction (Springer, 2009).

60. Li, S., Sesia, M., Romano, Y., Candès, E. & Sabatti, C. Searching for robust associations with a multi-environment knockoff filter. Biometrika 109, 611–629 (2022).

61. Heinze-Deml, C., Peters, J. & Meinshausen, N. Invariant causal prediction for nonlinear models. Journal of Causal Inference 6, 20170016 (2018).

62. Dixit, A. et al. Perturb-Seq: Dissecting Molecular Circuits with Scalable Single-Cell RNA Profiling of Pooled Genetic Screens. en. Cell 167, 1853–1866.e17 (Dec. 2016).

63. Wagner, A. et al. Metabolic modeling of single Th17 cells reveals regulators of autoimmunity. Cell 184, 4168–4185 (2021).

64. Cang, Z. et al. Screening cell–cell communication in spatial transcriptomics via collective optimal transport. Nature Methods 20, 218–228 (2023).

65. Vergara, H. M. et al. Whole-body integration of gene expression and single-cell morphology. Cell 184, 4819–4837 (2021).

66. Tung, P.-Y. et al. Batch effects and the effective design of single-cell gene expression studies. Scientific Reports 7, 39921 (2017).

67. Haghverdi, L., Lun, A. T., Morgan, M. D. & Marioni, J. C. Batch effects in single-cell RNA-sequencing data are corrected by matching mutual nearest neighbors. Nature Biotechnology 36, 421–427 (2018).

68. Gayoso, A. et al. A Python library for probabilistic analysis of single-cell omics data. Nature Biotechnology 40, 163–166 (2022).

69. Rozenblatt-Rosen, O., Stubbington, M. J., Regev, A. & Teichmann, S. A. The Human Cell Atlas: from vision to reality. Nature 550, 451–453 (2017).

70. Zou, H. & Hastie, T. Regularization and variable selection via the elastic net. en. Journal of the Royal Statistical Society 67, 301–320 (Apr. 2005).

71. Yao, Y., Rosasco, L. & Caponnetto, A. On Early Stopping in Gradient Descent Learning. en. Constructive Approximation 26, 289–315 (Apr. 2007).

72. Seabold, S. & Perktold, J. statsmodels: Econometric and statistical modeling with python in 9th Python in Science Conference (2010).

73. Stuart, T. et al. Comprehensive Integration of Single-Cell Data. en. Cell 177, 1888–1902.e21 (June 2019).

74. Petukhov, V. et al. Cell segmentation in imaging-based spatial transcriptomics. en. Nature Biotechnology 40, 345–354 (Mar. 2022).

75. FFPE Nanostring CosMX data https://nanostring.com/products/cosmx-spatial-molecular-imager/nsclc-ffpe-dataset/. Accessed: 2023-10-24.

